# Ecology, genomics and biocontrol potential of bacteriophages infecting the bacterial wilt pathogen *Ralstonia solanacearum* species complex in Réunion Island

**DOI:** 10.64898/2026.05.27.728151

**Authors:** Fernando Clavijo-Coppens, Sohini Claverie, Isabelle Robène, Stéphane Poussier, Laurène Robert, Maelle Pelissier, Julie Frapaise, Stéphanie Javegny, Murielle Hoareau, Claudine Boyer, Jean-Jacques Cheron, Jean-Michel Lett, Arthur Planche, Marie-Stéphanie Vernerey, Yann Pecrix, Adrien Rieux

## Abstract

The *Ralstonia solanacearum* species complex (RSSC), the causal agent of bacterial wilt, is among the most destructive soil-borne plant pathogens worldwide, yet effective and sustainable control strategies remain limited. Bacteriophages represent promising biocontrol agents, but their efficacy depends on ecological compatibility with local pathogen populations. Here, we combined ecological sampling, comparative genomics, phenotypic characterization and plant assays to investigate RSSC-infecting phages in Réunion Island and evaluate their biocontrol potential. We isolated 45 phages from diverse agricultural matrices and obtained complete genome sequences for 35 novel isolates. Phylogenomic analyses revealed a locally diversified assemblage comprising multiple known taxa and several putative new genera, forming clusters largely distinct from global reference phages. Phage diversity and antibacterial activity were structured primarily by bacterial phylogeny rather than plant host or geographic origin, indicating that plants act mainly as ecological interfaces while environmental bacterial populations shape phage specialization. The community displayed two contrasting evolutionary strategies: expanding virulent lineages associated with strong antibacterial activity and persistent temperate lineages carrying integration and host-interaction functions. Host-range assays confirmed phylotype-dependent susceptibility, and strictly lytic phages showed consistently higher inhibitory activity. Guided by combined genomic and phenotypic screening, we designed a multi-family phage cocktail targeting dominant local RSSC lineages. The cocktail exhibited strong *in vitro* suppression of bacterial growth and significantly reduced disease severity in tomato plants. Together, our results demonstrate that effective phage biocontrol depends on evolutionary matching between phages and regional pathogen populations. Integrating ecological, genomic and functional characterization provides a robust framework for selecting locally adapted phages and developing durable phage-based strategies for managing bacterial wilt.

## Introduction

The *Ralstonia solanacearum* species complex (RSSC), the causal agent of bacterial wilt, infects more than 200 plant species and is considered one of the most destructive plant pathogens worldwide, particularly in tropical and subtropical regions where warm and humid conditions favor its proliferation and dissemination (Mansfield et al. 2012). Owing to its extensive genetic and phenotypic diversity, RSSC is classified as a species complex comprising three species (*R. solanacearum*, *R. pseudosolanacearum* and *R. syzygii*) and four major phylogenetic groups known as phylotypes (Prior et al. 2016). In Réunion Island, RSSC severely affects economically important crops such as potato, tomato, eggplant and pepper, posing a major threat to agricultural sustainability and food security. Local surveys have revealed the predominance of *R. pseudosolanacearum* phylotype I (Yahiaoui et al. 2017; Cellier et al. 2023). Control of RSSC remains particularly challenging because the bacterium persists in soil and water and rapidly recolonizes cultivated fields despite crop rotation or sanitation practices (Hayward 1991). Chemical options are limited and resistant cultivars remain scarce, highlighting the need for sustainable management strategies (Elphinstone, J. 2005).

Bacteriophages (phages), viruses infecting bacteria, have emerged as promising biocontrol agents offering targeted and environmentally friendly alternatives to chemical pesticides (Buttimer et al. 2017; Holtappels et al. 2021). Their potential has been demonstrated against several plant pathogens, including *Pseudomonas syringae*, *Xanthomonas* spp., *Xylella fastidiosa* and *Erwinia amylovora* (Clavijo-Coppens et al. 2022). In RSSC, phage cocktails have successfully reduced disease incidence in tomatoes under greenhouse and field conditions (Bae et al. 2012; Álvarez et al. 2019; Wang et al. 2019). However, treatment efficacy remains variable and often poorly reproducible, suggesting that selecting phages solely based on lytic activity or host range may be insufficient (Holtappels et al. 2021).

Recent advances in viral ecology indicate that phage–host interactions are strongly constrained by bacterial population structure and evolutionary history rather than random encounter alone (Roux et al. 2016; Pfeifer et Rocha 2024; Pfeifer et al. 2022). Phylogeny-informed host-range analyses show that bacterial genetic diversity shapes phage specialization and virulence, with implications for therapy durability (Torres-Barceló et al. 2025; Feltin et al. 2025). Moreover, resistance evolution and ecological trade-offs can determine long-term stability of phage treatments (Baud et al. 2025), while rhizosphere phage communities may influence disease outcomes at the ecosystem level (Yang et al. 2023). Together, these findings suggest that phage biocontrol success depends not only on phage identity but on its evolutionary compatibility with local pathogen populations. Yet this relationship remains poorly tested *in planta* pathosystems, where phage selection is still largely empirical.

Here, we investigated whether the ecological and evolutionary structure of local RSSC populations determines the diversity and efficacy of associated bacteriophages. Building on previous work in the Southwest Indian Ocean (Trotereau et al. 2021), we combined environmental sampling, comparative genomics and phenotypic assays to characterize RSSC phages in Réunion Island. We then used these data to design a locally adapted phage cocktail and evaluated its performance *in planta*. By linking phage ecology, host population structure and biocontrol outcome, this study aims to provide a framework for rational selection of phages for sustainable bacterial wilt management.

## Methods

### Sampling, isolation and purification of phages

Sampling was carried out between 2018 and 2021 across eight agricultural plots located on the west coast of Réunion Island (**Figure S1**). Sampled sites were identified based on suspected cases of bacterial wilt in solanaceous crops, regardless of geographic location, host plant species, or farming system (e.g., greenhouse, open field, soilless, or organic cultivation). All samples consisted of soil collected from around plants (potato, eggplant, pepper tomato & datura) roots, except for one sample, which was composed of irrigation water. In the laboratory, samples were enriched overnight using a liquid culture of strain RUN3012 (sequevar I-31), the most commonly reported sequevar in Réunion Island. Phages were isolated using the overlay plaque assay method. Individual plaques were picked and purified through three rounds of re-plating. Phage lysates were then prepared using a standard chloroform-based protocol, and their concentration was determined by counting plaque-forming units (PFU) at the highest dilution. Final phage solutions were kept at 4 °C until use.

### Bacterial strains and culture media

All bacterial strains used in this study are part of the reference collection at the Pôle de Protection des Plantes in Réunion Island, France. **Table S1** provides details on their phylogenetic classification and origin. All inoculations and bacterial cultures of RSSC strains were performed using a semi-selective Kelman medium supplemented with 1% m/m triphenyltetrazolium chloride, whereas non-RSSC bacterial strains were cultured on LPGA medium. Bacterial cultures were incubated at 28°C, with liquid cultures agitated at 80 rpm in the presence of phages or 120 rpm for bacteria alone.

### In-vitro measurement of phage host range and virulence

Virulence (the efficiency with which a phage infects, replicates, and lyses bacterial cells) was quantitatively measured using the bacterial growth kinetics method in a Bioscreen 100-well plate. In brief, an overnight culture of RUN3012 strain was diluted to an OD600 of 0.1 in fresh broth to ensure log-phase growth during the experiment. Following concentration normalization, phages were either added individually or in cocktail to RUN3012 culture at ratio 1:100 in a 300 µL final volume. When tested in cocktails, phages were mixed in equal quantities. OD600 measurements were taken every 15 minutes during 24 hours at 28°C with continuous shaking. Control wells contained bacteria without phages and a blank medium control. Each condition was tested in triplicate to ensure reproducibility. At the end of the experiment, virulence was calculated using the following formula:

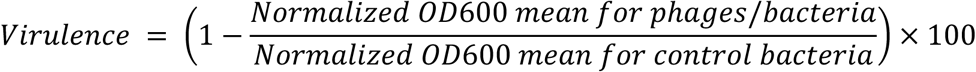

Hence, a virulence of 100% indicates a total inhibition of bacterial growth by phages while a virulence of 0% designates no inhibition of bacterial growth by phages at all, meaning an unaffected bacterial growth, as compared to the control.

Host range (the spectrum of bacterial strains that a phage can infect) measurement was performed using the spot assay. Each tested phage was spotted at various concentrations (ranging from 10^9^ to 10^6^ PFU.mL^-1^) onto a selection of bacterial strain lawns. After a 24-hour incubation period, the presence of plaques indicates that the phage successfully infected the bacterium, whereas their absence suggests the phage was unable to infect it.

### Phage DNA extraction, sequencing and assembly

Genomic DNA extraction was performed with the Wizard DNA clean-up kit from Promega. DNA was converted into a double-stranded library either using theTruSeq DNA Nano Illumina protocol or the Nanopore rapid Barcoding Kit 24. Sequencing was either performed in a paired-end 2×150 cycles configuration on a single lane of the HiSeq flow cell or in a single-end mode on a MinION flow cell, respectively. Adaptors and quality trimming was done with Trimmomatic 0.36 (Bolger et al. 2014). Trimmed Illumina-only, Nanopore-only or Illumina+Nanopore reads were *de novo* assembled using Unicycler v0.5.0 (Wick et al. 2017) with default parameters. A total of 45 annotated Ralstonia phage genomes were submitted to the European Nucleotide Archive (ENA) under BioProject accession PRJEB111082. Each genome is associated with a dedicated BioSample accession and ENA analysis accession as indicated in **Table S2**.

### Genomic analyses

Genome assembly quality was evaluated using Quast (Gurevich et al. 2013), including evaluation of contiguity, genome size, GC content, and coverage metrics. Genome annotation was performed using a complementary pipeline combining RAST®, PHASTEST, and VirClust in order to improve functional prediction and taxonomic contextualization (Wishart et al. 2023; Moraru 2023; Aziz et al. 2008). Coding sequences (CDSs) were predicted automatically by these three tools, and functional annotations were curated based on conserved domains and homology searches using BLASTp and HHpred blast databases. Phage lifestyles, classified as either lytic or lysogenic, were inferred based on conserved protein domains using Bacphlip (Hockenberry et Wilke 2021). To assess genomic relatedness and facilitate taxonomic classification at the family, genus, and species levels, nucleotide-based intergenomic similarity indices were calculated using Viridic & Victor following ICTV-recommended thresholds for species and genus demarcation (Moraru et al. 2020; Meier-Kolthoff et Göker 2017). In addition, genome-wide phylogenetic relationships were further explored using the alignment-free method implemented in Mashtree to confirm clustering robustness independently of alignment-based reconstruction (Katz et al. 2019). Global genome relatedness was further evaluated using the shared *k*-mer approach implemented in the KI-S (K-mer–based Identification of Similarity) framework (Briand et al. 2021). Pairwise similarity matrices were computed using the KI-S tool available on the CIRM-CFBP Galaxy platform (https://iris.angers.inra.fr/galaxypub-cfbp/). The resulting similarity matrix was visualized using the KI-S circle packing representation, providing a non-hierarchical projection of genome similarity relationships that complements tree-based phylogenomic analyses. To contextualize the genetic diversity of RSSC-phages Réunion Island, we compared them against a curated dataset of complete *Caudovirales phages* genomes infecting members of the order *Burkholderiales* available in GenBank (**Table S3**).

### Transmission electron microscopy (TEM) photographs

Transmission electron microscopy was performed at the PHIM imaging platform (Plant Health Institute of Montpellier, France). Briefly, 2 mL of phage suspension were centrifuged at 20,800×g for 1 h at 4 °C. The resulting pellet was washed three times with 2 mL of TEM buffer (0.1 M ammonium acetate, pH 7, 0.22 µm-filtered) and resuspended in a final volume of 50 µL. Aliquots of 5 µL (∼2 × 10^8 PFU·mL−1) were deposited onto glow-discharged carbon-coated copper grids (EMS-D300FC-50Cu) and allowed to adsorb for 5 min. Grids were negatively stained with drops of 2%(w/v) of uranyl acetate for 3 min. After drying on filter paper, samples were rinsed on 3 drops of water and then examined with a JEOL JEM-1400 Plus transmission electron microscope. Images were further processed with ImageJ software.

### Evaluation of phage cocktail efficacy under in planta conditions

Tomato plants (*Solanum lycopersicum* cv. Roma) were germinated at 28 °C under a photoperiod of 16h light / 8h dark and transplanted at two weeks into pots containing 100 g of commercial substrate. Bacterial infection was performed by mechanically wounding the roots with a sterile scalpel, followed by watering with 50 mL of a RUN3012 strain suspension (2 × 10⁷ CFU·mL⁻¹, corresponding to approximately 10⁹ CFU per plant). Five treatments were evaluated using ten plants per condition and three biological replicates: (i) negative control (uninfected plants + no phage treatment), (ii) phage-cocktail control (applied as irrigation water on uninfected plants throughout the experiment), (iii) bacterial infection control (no phage treatment), (iv) single phage cocktail application, and (v) continuous phage cocktail application. For the single application, substrates were drenched with 50 mL of phage cocktail (2 × 10⁶ PFU·mL⁻¹; approximately 10⁸ PFU per plant or 10⁶ PFU·g⁻¹ of substrate equivalent to a phage-to-bacteria ratio of 1:10) two days prior to bacterial inoculation. The continuous treatment began in the same way, followed by twice-weekly phage applications thereafter. Disease severity was scored on a 0–4 scale: 0 = healthy, 1 = one symptomatic leaf, 2 = <50% wilt, 3 = >50% wilt, 4 = complete wilt or plant death, as previously described (Kempe 1983).

## Results

### Phages isolation, sequencing and assembly

Agricultural samples were collected across contrasting agroecosystems of Réunion Island and phage-enriched with strain RUN3012 to capture the diversity of Ralstonia-associated bacteriophages associated with RSSC-infected crops. A total of 45 *Ralstonia*-associated phage isolates were recovered and purified from agricultural samples, predominantly soils (97%) and to a lesser extent irrigation water (3%). These samples were collected from greenhouses (60%) and open fields (40%) associated with RSSC-infected datura (5%), eggplant (20%), tomato (63%), potato (5%), and pepper (7%) plants located both in the southern (75%) and western (25%) part of Réunion Island (**Table 1 & Figure S1)**. After purification, phage genomic DNA was extracted and individual phage isolates were subjected to whole-genome sequencing. Among these, ten of our phage isolates had been previously characterized in an earlier study (Trotereau et al. 2021). Genome assembly of the remaining 35 phages resulted in complete, single-contig genomes ranging from 41 to 91 kb in length, with GC contents between 53% and 64% and comprising 44 to 80 predicted coding sequences (**Table 1**).

**Table 1.**
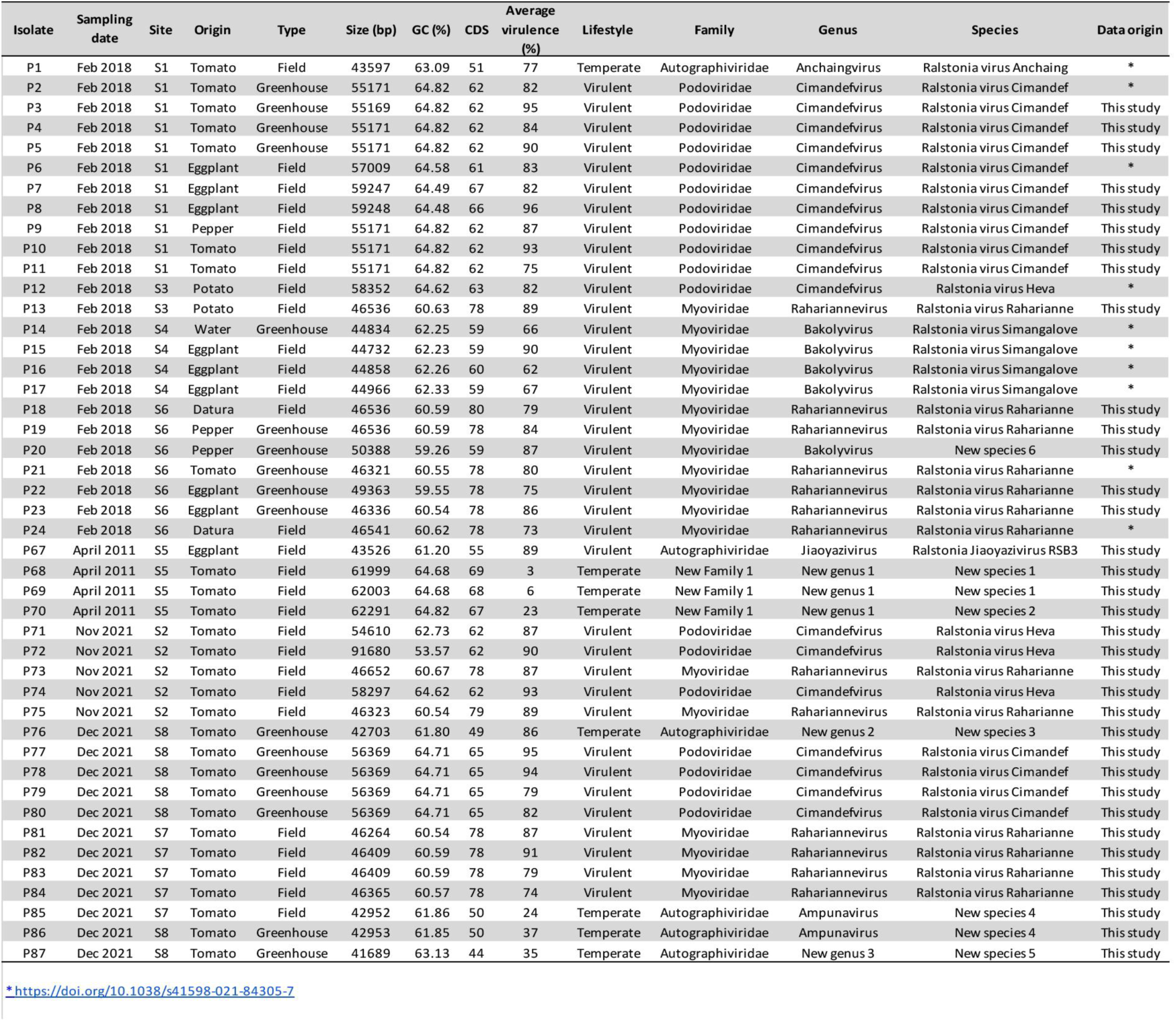
Genomic, ecological, and taxonomic characteristics of the 45 bacteriophages isolated in this study.

### Individual phage life-history traits assessment and functional genomic architecture

*In-vitro* phage virulence was assessed through bacterial growth kinetics. Control cultures of the RUN3012 strain exhibited a standard log-phase growth curve, with no contamination detected in blank medium controls. Among the 45 tested phages, virulence varied from 3% (P68) to 95% (P8), with 36 phages demonstrating virulence above 70% (**Table 1**). As expected, phage lifestyle was strongly associated with virulence: lytic phages consistently exhibited high inhibition levels against strain RUN3012 (>80%), whereas temperate phages showed lower and more variable inhibitory activity (<40%). Importantly, no growth-promoting or antagonistic effects were detected, nor were any bacteriostatic profiles observed, indicating that differences in virulence reflect robust lytic activity rather than transient growth arrest. *In-silico* genomic analysis performed with BACPHLIP classified 8 phages as temperate (lysogenic) and 37 as virulent (lytic) (**Table 1**).

Genome annotation revealed a non-uniform distribution of functional gene categories across the 45 *Ralstonia solanacearum*–infecting phages, strongly structured by phage lifestyle **(Table 2)**. Integrase and excisionase genes were detected exclusively in temperate phages: classical lysogeny-associated modules, combining a site-specific tyrosine recombinase-type integrase and an excisionase, were identified in phages P68, P69 and P70, while P1, P85 and P86 carried integrase genes only (**Table S4**). In contrast, no integrase nor excisionase was detected in strictly virulent phages. Conversely, recombination-related functions unrelated to lysogeny were widespread among lytic phages, which frequently encoded transposases, including homologues related to CRISPR-Cas transposon-associated proteins (COG0675: InsQ/TnpB), notably in lineages P2–P12, P18–P24, P71–P75 and P77–P84, as well as shufflon-specific DNA recombinases in phages P76 and P87. Beyond recombination functions, adaptive genes were prevalent across the phage collection: 39 phages encoded at least one auxiliary metabolic gene (AMG), and 26 carried host interaction-related genes. AMGs linked to plant-associated functions, such as polysaccharide degradation and lipid metabolism, were predominantly detected in virulent phages of the genera Cimandefvirus and Rahariannevirus (e.g. pectin lyase-like proteins in P2–P12, P71–P75; esterase/lipase genes in P18–P24, P73 and P75). In contrast, temperate phages more frequently encoded AMGs related to nucleotide metabolism and iron acquisition, including deoxynucleoside monophosphate kinase, phosphoribosyl transferase and Iron(III) dicitrate transporters (P76, P85 and P86), as well as host interaction genes such as SAM-dependent methyltransferases, restricted to phages P68–P70. In addition, several phages, including P1, P76, P85, P86 and P87, encoded a phage-specific DNA-directed RNA polymerase, indicating partial transcriptional autonomy from the host machinery. Hypothetical proteins represented the largest category in all genomes (35–62%), underscoring the extensive uncharacterized functional diversity of RSSC-associated phages.

**Table 2.**
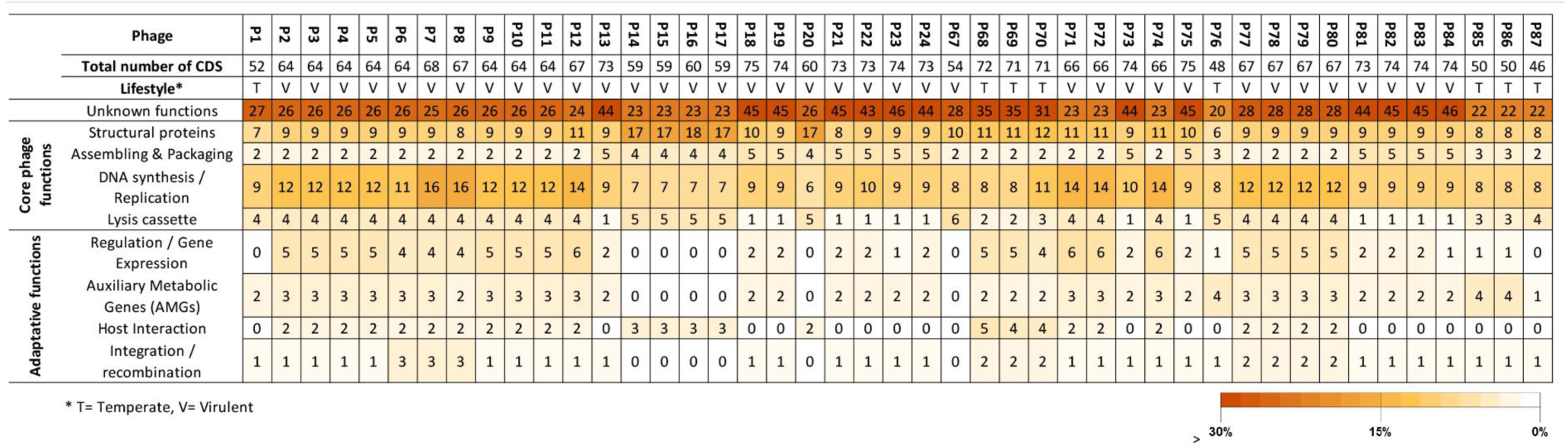
Annotation of the 45 Ralstonia solanacearum–infecting phage genomes. Values indicate the number of CDS per genome assigned to each functional category, while colors represent the corresponding percentage. Core phage functions, adaptive functions and regulatory or hypothetical genes are distinguished.

### Taxonomic assignment

Genome-wide taxonomic assignment of the 45 bacteriophages isolated in Réunion Island was performed using a combination of *BLASTn* nucleotide similarity searches, genome size and GC content comparisons, following ICTV recommendations. These analyses revealed that a large fraction of the phages showed very high nucleotide similarity (>99% identity over >94% genome coverage) to previously described *Ralstonia solanacearum* phages, allowing their assignment to recognized species, including *Ralstonia virus Anchaing, Cimandef, Heva, Raharianne, and Simangalove*. At the genus level, these phages were distributed across four recognized genera: Anchaingvirus and Jiaoyazivirus (Autographiviridae), Cimandefvirus (Podoviridae), and Bakolyvirus and Rahariannevirus (Myoviridae). Phage P67 showed its closest relationship to Ralstonia phage RSB3, sharing 95.89% nucleotide identity over 97% genome coverage, supporting its assignment to the genus *Jiaoyazivirus*. Similarly, phages P85 and P86 exhibited their highest similarity to *Burkholderia* phage AMP1, sharing 85.5–87% nucleotide identity over 87–89% genome coverage, supporting their inclusion within the genus Ampunavirus (Autographiviridae), while representing a distinct putative novel species according to ICTV species demarcation criteria (i.e. below the 95% nucleotide identity threshold). In contrast, several phages displayed lower nucleotide similarity to any previously described genomes. Phage P20 clustered within the genus *Bakolyvirus* but was clearly separated from known species, supporting its classification as a novel *Bakolyvirus* species. Phages P68, P69, and P70 showed limited similarity (<40% genome coverage; ∼80–81% nucleotide identity) to *Burkholderia* phage JC1 and were classified as two new species within a putative novel genus belonging to a previously undescribed family. Phage P76 showed partial similarity to *Burkholderia* phage AMP3 (87% genome coverage, 83.8% nucleotide identity) supporting the proposal of a second putative novel genus within *Autographiviridae*. Finally, phage P87 exhibited limited similarity to *Ralstonia* phage RS-PII-1 (77% genome coverage, 82% nucleotide identity), consistent with the delineation of a third putative novel genus and species. Moreover, P1, P76, P85, P86 and P87 encoded a phage-specific DNA-directed RNA polymerase, a hallmark of the *Autographiviridae* family, supporting their assignment to this taxon. All together, taxonomic analyses supported the delineation of eleven species distributed across nine genera, including three putative novel genera and several novel species, within the collection of RSSC-associated phages isolated in Réunion Island (**Table 1**).

### Phylogenetic diversity at both local and global scales

The phylogenetic tree built with Mashtree from the 45 Réunion phage genomes resolved into three major clades (**Figure 1A)**. Clade 1, located at the base of the tree, comprises all members of the *Autographiviridae* family, including *Jiaoyazivirus, Ampunavirus, Anchaingvirus* and two newly identified genera. This clade displays substantial genomic divergence, consistent with the high evolutionary plasticity reported for members of this family. Clade 2 consists exclusively of temperate phages, all assigned to a putative newly identified family. Clade 3 encompasses both *Podoviridae,* including multiple *Cimandefvirus* and *Anchaingvirus* members forming a cohesive and genomically homogeneous clade, and *Myoviridae*, represented by *Bakolyvirus* and *Rahariannevirus*, which constitute distinct yet related sublineages within this group. Overall, the observed phylogenetic structure was entirely congruent with the taxonomic classification, demonstrating a strong alignment between genomic similarity and taxonomic structure. In contrast, phage diversity did not appear to be strongly structured by spatial location, crop identity or type of production system. These results were further supported by another phylogenomic inference based on intergenomic distances calculated using VICTOR (nucleotide-based, D0 formula, trimming), which produced highly supported topologies (branch support >90%) fully consistent with the mashtree reconstruction (**Figure S2**).

**Figure 1.**
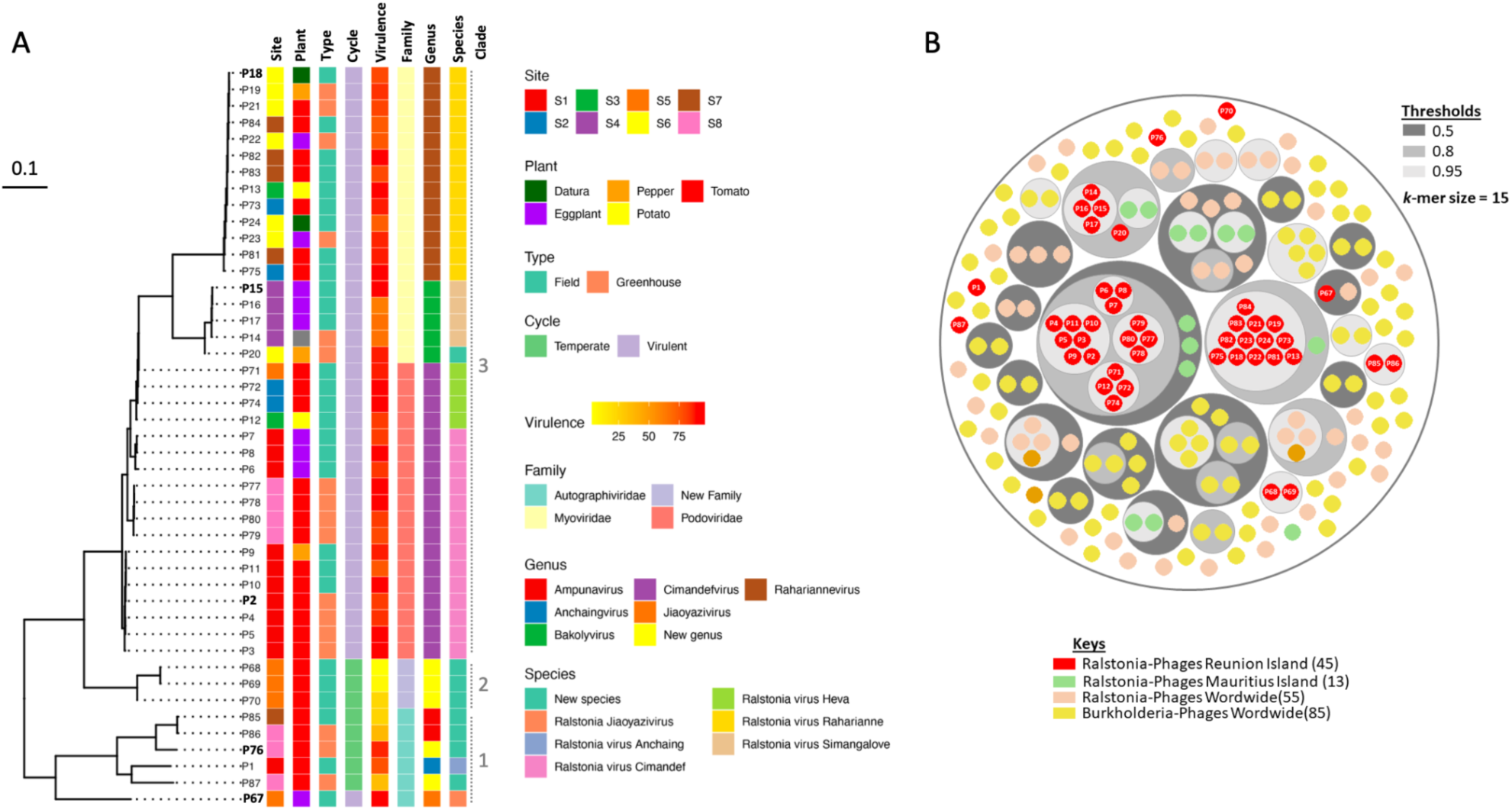
Local diversity and global positioning of Ralstonia phages isolated on Réunion Island. (A) Phylogenetic tree of the 45 *Ralstonia*-infecting bacteriophages isolated on Réunion Island, reconstructed from whole-genome comparisons. Branch lengths represent Mash distances and phages selected for the cocktail are shown in bold. The accompanying heat map summarizes sampling information together with inferred lifestyle, virulence, and taxonomic classification. (B) KI-S circular plot illustrating genome relatedness between Réunion Island phages (red dots) and publicly available *Ralstonia* and *Burkholderia* phages worldwide. Concentric circles represent genome clusters defined at three similarity thresholds (0.5, 0.8, and 0.95; grey gradients), based on k-mers of size 15. Colors denote the geographic origin of the phages, and numbers correspond to the IDs of Réunion Island phages only.

Genome relatedness analysis based on KI-S projection was used to position the 45 bacteriophages isolated in Réunion Island within the global diversity of Ralstonia- (68 genomes) and Burkholderia-infecting (85 genomes) phages (**Figure 1B**). In this multidimensional genomic space, all 45 Réunion Island phages grouped within confined clusters of the projection, clearly separated from the majority of reference phages isolated outside the Southwest Indian Ocean region. No Réunion Island phages overlapped with distant continental clusters, indicating a marked geographic structuration of genome relatedness. Within this region, Réunion Island phages were structured into multiple groups fully congruent with their taxonomic assignment. The closest genetic affinities were systematically observed between phages isolated in Réunion Island and those originating from the neighboring island of Mauritius, supporting a strong geographic signal in phage genome relatedness. With the notable exception of phage P67, which grouped with Ralstonia phage RSB3, for which no geographic origin is reported, no Réunion Island phages showed close genomic proximity to phages isolated outside the Southwest Indian Ocean region. Importantly, reference phages from other continents occupied distinct and largely non-overlapping regions of the KI-S projection. Together, these results indicate that while Ralstonia- and Burkholderia-infecting phages exhibit extensive global genomic diversity, Réunion Island represents a localized hotspot of phage diversification area within this broader genomic landscape.

### Host-range assessment and phage selection for cocktail development

Based on *in vitro* inhibitory activity (>80%), a strictly virulent lifestyle, phylogenetic distinctiveness, absence of toxin- or virulence-coding genes, and the absence of negative interactions beyond the reference strain RUN3012, ten phages (P3, P7, P8, P15, P18, P19, P67 & P82) were selected for host range evaluation. This subset included representatives of five genera across three families, with deliberate redundancy within the dominant genus Cimandefvirus (6/10 phages) to assess intra-genus variability in host range patterns. These phages were tested against 25 RSSC strains representing phylotypes I, II, III, and IV from the Indian Ocean region, as well as nine non-host bacterial strains isolated from other regions (Table S1, **Figure 2**). Overall, phage activity was not associated with the geographic origin of bacterial isolates but was instead strongly structured by bacterial phylotype. All 10 phages exhibited lytic activity against at least one phylotype I strain, encompassing sub-lineages I-13, I-18, and I-31 originating from La Réunion, Mauritius, and Madagascar. In contrast, no lytic activity was detected against phylotype I strains belonging to sequevars I-15, I-33, and I-46, regardless of their geographic origin. Activity against phylotype II strains was more limited: only phages P2 and P8 were active against both II-B-1 strains from La Réunion and Madagascar, whereas P71 was active against the Madagascar isolate only. No lytic activity was detected against any phylotype III or IV strains, regardless of geographic origin. Importantly, none of the tested phages exhibited lytic activity against non-host bacteria, including *Pseudomonas* spp., *Escherichia coli*, *Salmonella typhi*, *Bacillus subtilis*, or non-RSSC *Ralstonia* species, confirming their narrow host specificity. Based on these results, five phages (P2, P15, P18, P67, and P76), representing three families and five genera, were selected to prepare a phage cocktail designed to ensure the broad coverage of local *R. solanacearum* strain diversity.

**Figure 2.**
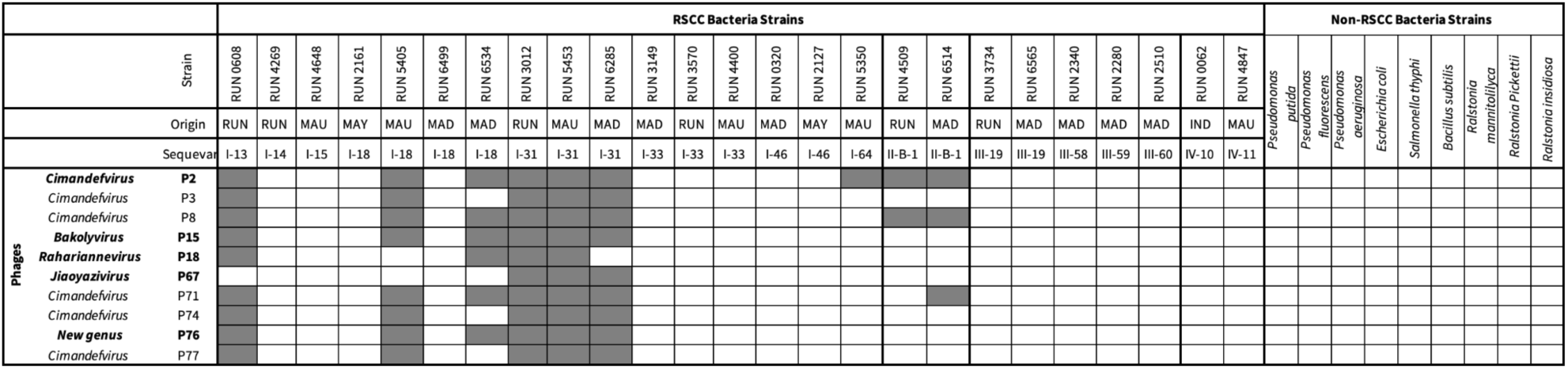
Host range of selected bacteriophages on Ralstonia isolates and non-host bacteria. Binary heatmap showing the lytic activity of selected phages against *Ralstonia solanacearum* species complex isolates representing multiple phylotypes, sequence variants and geographic origins, including Réunion Island (RUN), Mauritius (MAU), Madagascar (MAD), Mayotte (MAY) and Indonesia (IND). Phages are grouped by genus. Grey cells indicate lysis. Non-RSSC strains were included as specificity controls. Phages in bold were selected for cocktail formulation.

### Phage morphology and in vitro combination assays

Transmission electron microscopy revealed distinct morphotypes among the five selected phages used for cocktail development (**Figure 3**). Phages P2, P67, and P76 exhibited isometric capsids, with diameters of approximately ∼54.36 nm, ∼62.81 nm, and ∼80.82 nm, respectively. Based on capsid morphology and tail structure, P2 displayed features consistent with a podophage morphotype, whereas P67 and P76 exhibited morphologies consistent with *Autographiviridae*-like phages. Short tails were observed, but their small size precluded accurate measurements. In contrast, phages P15 and P18 displayed morphologies characteristic of myophages, with long flexible tails attached to isometric capsids. Phage P15 exhibited an isometric head of approximately ∼44.94 nm in diameter and a tail length of approximately ∼102.39 nm. Phage P18 showed a similar overall morphology, with an isometric capsid measuring approximately ∼55.96 nm in diameter and a tail length of approximately ∼94.22 nm.

**Figure 3:**
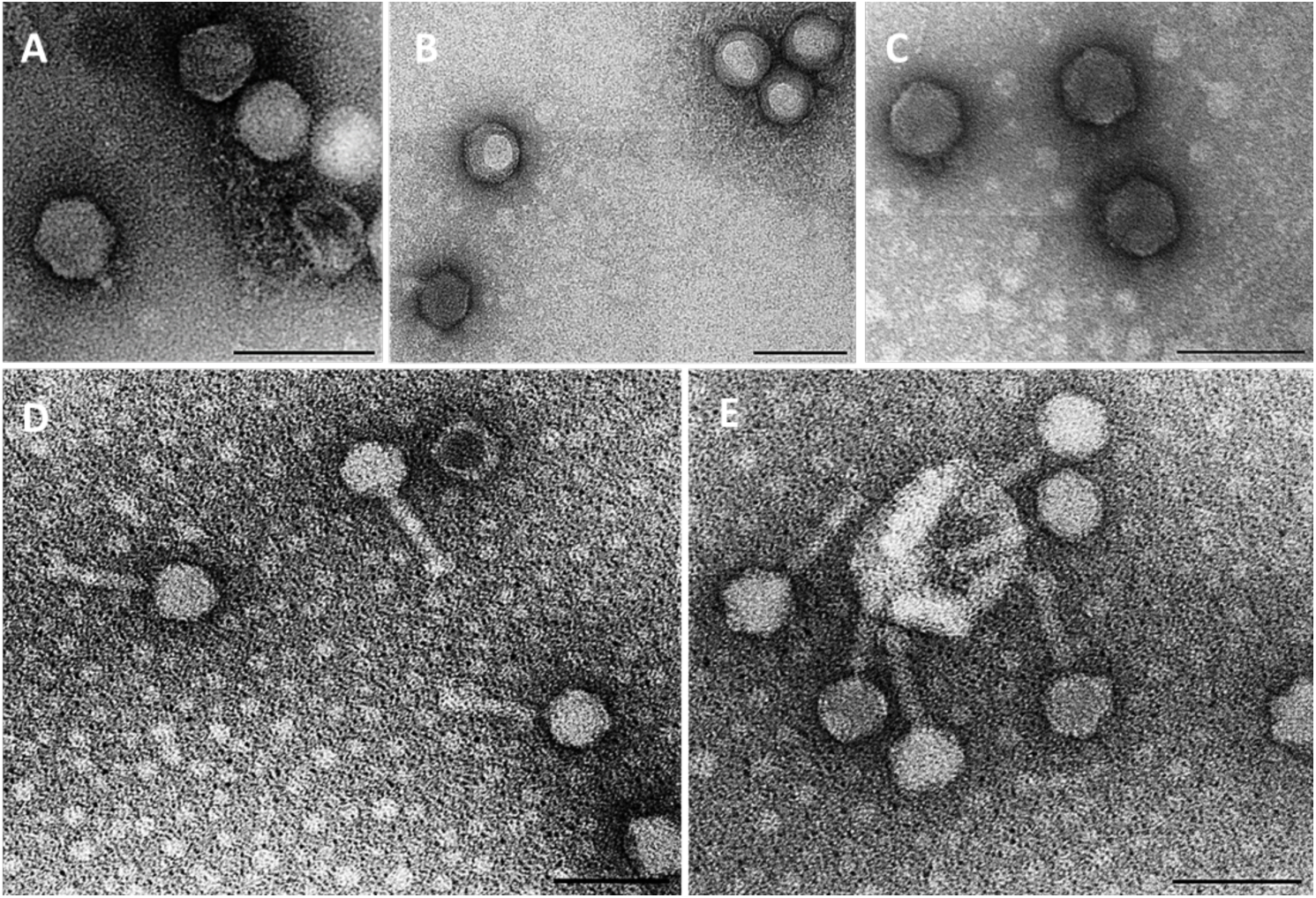
Transmission electron microscopy of five representative phages revealed distinct morphotypes. P2, P67, and P76 exhibited isometric capsids with short tails, consistent with podophage- and Autographiviridae-like morphologies (A-C). In contrast, P15 and P18 displayed typical myophage morphologies, with isometric capsids and long flexible tails (D-E). Scale bars= 100 nm are shown.

When tested *in vitro*, the phage cocktail induced strong and sustained suppression of bacterial growth, indicating robustness, lytic activity and the absence of antagonistic interactions among the constituent phages (**Figure 4**). No negative synergy was observed, as the virulence of phages mixture did not significantly decrease compared to individual phage applications. Relative virulence values ranged from 0.76 to 0.87, across all treatments, reflecting substantial inhibition of bacterial growth. A clear trend toward increased virulence was observed with increasing cocktail complexity. Median virulence values increased from approximately 0.80 for single-phage treatments to 0.83–0.84 for two-phage combinations, ∼0.85 for three-phage combinations, and 0.86–0.87 for four-phage combinations (**Figure S3**). The five-phage cocktail displayed one of the highest inhibitory activities (virulence ≈0.86), comparable to the upper range observed for four-phage combinations. This suggests that the different phages act in a non-antagonistic and potentially complementary manner, maintaining effective lytic activity against the target bacterial population when combined.

**Figure 4:**
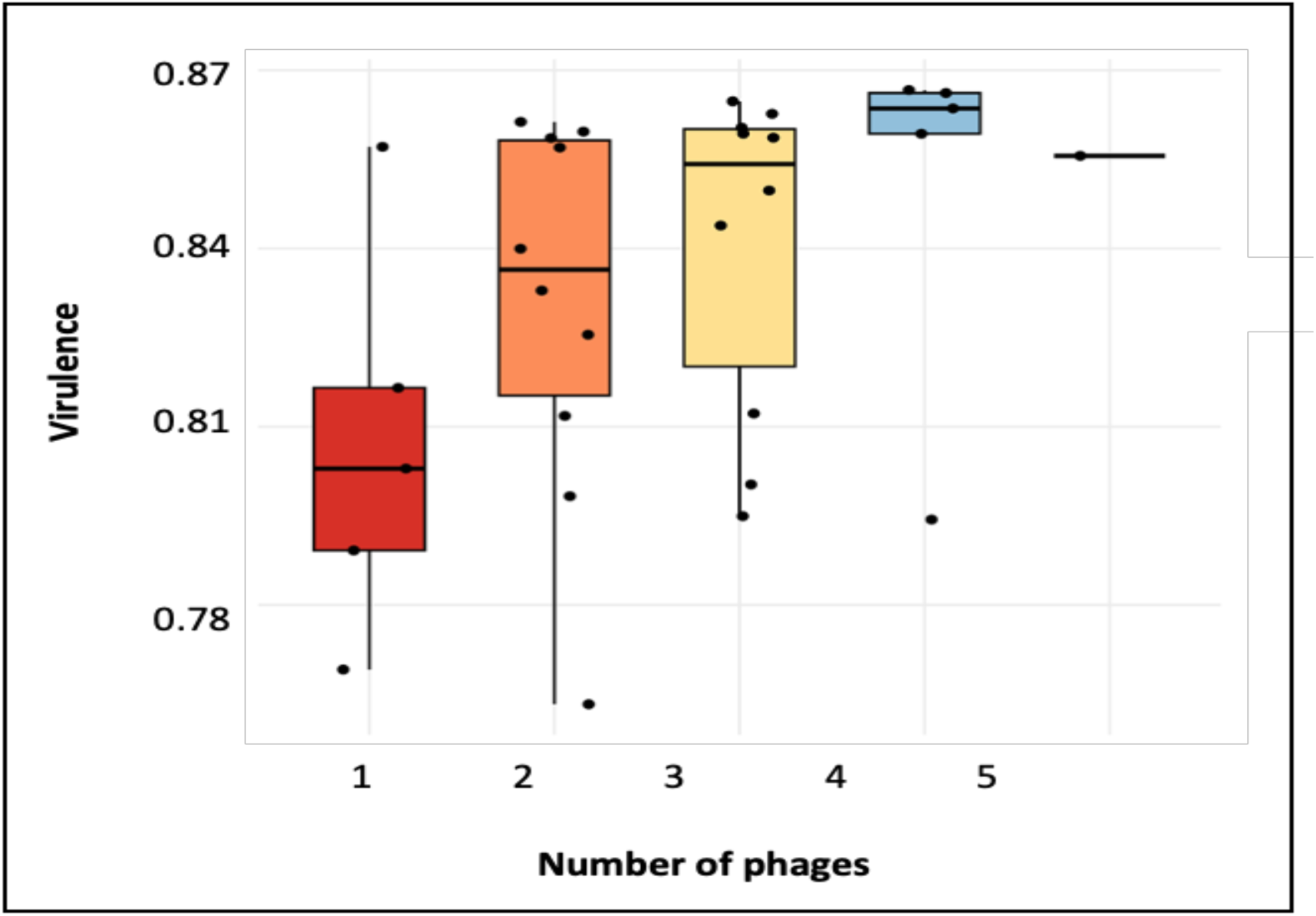
Comparison of *in-vitro* measured virulence among treatments containing 1, 2, 3, 4, or 5 phages. Boxplots show the median, interquartile range, and individual data points.

### In planta testing of the phage cocktail

The efficacy of the phage cocktail was evaluated on experimentally infected tomato plants under controlled conditions. Temporal analysis revealed clear differences in disease progression among treatments (**Figure 5A**). Infected control plants showed a rapid increase in symptom severity, reaching mean disease scores close to the maximum value by nine days post infection. In contrast, plants treated with the phage cocktail exhibited delayed symptom onset and reduced disease progression relative to infected controls. This effect was observed for both treatment modalities, with continuous phage irrigation consistently resulting in consistently lower mean symptom scores than a single application. Quantification of disease severity using the area under the disease progress curve (AUDPC) confirmed these observations quantitatively (**Figure 5B**). Infected control plants displayed the highest AUDPC values, whereas both phage-treated groups showed markedly reduced disease severity. Continuous phage application resulted in the lowest AUDPC values, followed by the single-application treatment, while uninfected control plants remained symptom-free throughout the experiment. Representative images further illustrated these differences in disease outcome among treatments (**Figure 5C**). After 20 days of monitoring, plants treated with continuous phage irrigation exhibited a survival rate of 53.3%, compared with 36.7% for plants receiving a single phage application. In contrast, all untreated infected control plants succumbed to the disease (0% survival). Uninfected control plants, irrigated with either phage buffer or water, maintained 100% survival, indicating that phage application was not phytotoxic under the tested conditions.

**Figure 5:**
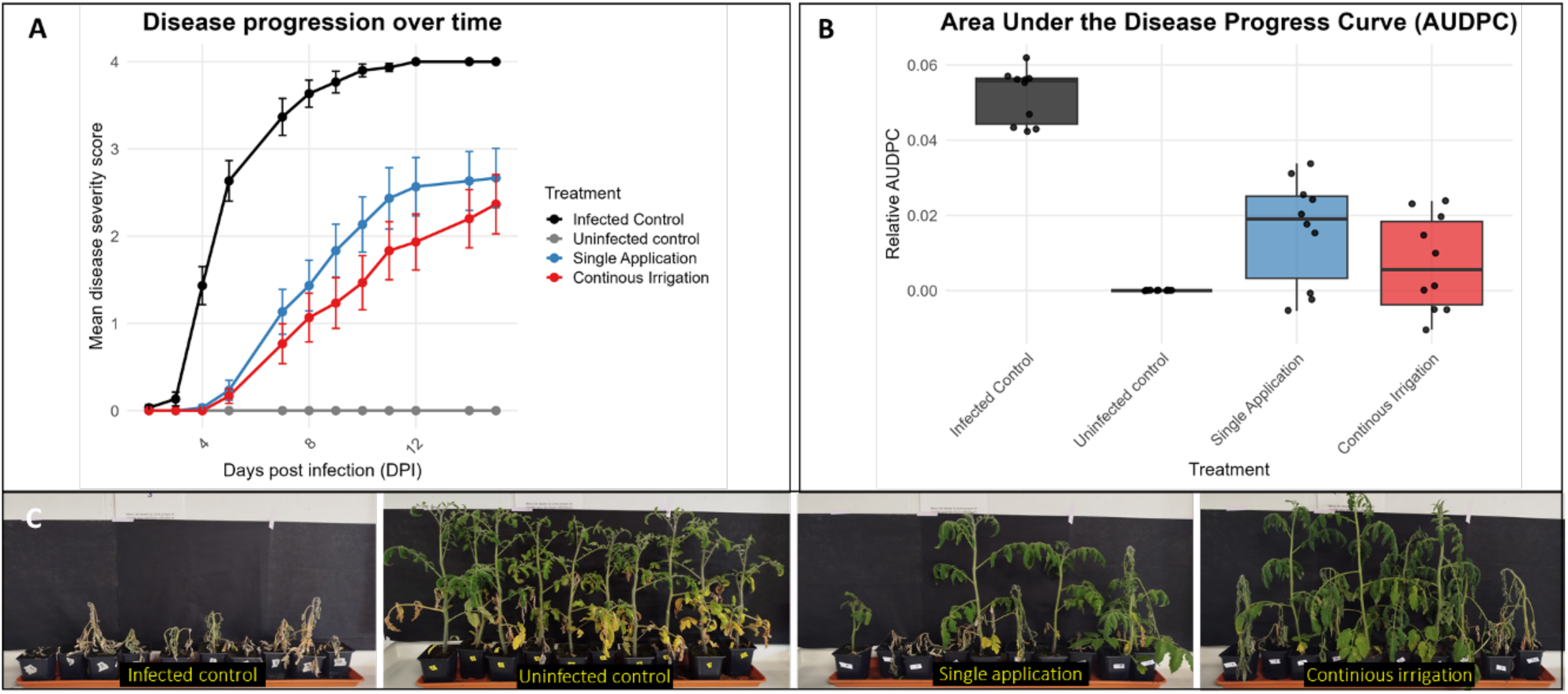
*In planta* evaluation of a five-phage cocktail against bacterial wilt of tomato. (A) Mean disease severity scores (0–4 scale) of tomato plants infected with *Ralstonia solanacearum* strain RUN3012 under different treatments over 15 days post-inoculation (dpi). (B) Area under the disease progress curve (AUDPC) summarizing disease severity over time. (C) Representative images of tomato plants 20 days after inoculation. Left panel: infected control; second panel: uninfected control; third panel: single phage application; right panel: continuous phage irrigation.

## Discussion

### RSSC phages form an environmental reservoir structured by host populations rather than plant environments

This study provides the most comprehensive characterization to date of bacteriophages associated with the RSSC in Réunion Island. The recovery of 45 phages across multiple agricultural matrices demonstrates that RSSC phages are widespread throughout agroecosystems affected by bacterial wilt. Although sampling was not designed to resolve spatial distribution, their detection in all surveyed environments supports the existence of a broadly distributed environmental reservoir, as previously suggested in various bacterial wilt regions (Trotereau et al. 2021; Álvarez et al. 2019; Wang et al. 2019). Despite ecological heterogeneity among sampling sites, crop species, and production systems, neither phage diversity nor antibacterial activity showed consistent association with plant origin. Instead, host range was primarily determined by bacterial phylotype. This indicates that plants act mainly as ecological interfaces where interactions become observable rather than selective environments driving phage specialization. Phage population structure therefore appears to be shaped predominantly by shared environmental bacterial reservoirs rather than by individual agroecosystems. Beyond phylogenetic diversity, phages also displayed substantial functional variation, including genes involved in host metabolism and host–phage interactions, indicating diversification at both genetic and ecological levels.

### Coexistence of persistent temperate lineages and expanding virulent lineages

The RSSC phage community exhibited two contrasting evolutionary patterns. Strictly lytic lineages, particularly within *Cimandefvirus* and *Rahariannevirus*, consisted of closely related isolates consistent with recent local expansion, whereas temperate phages formed phylogenetically isolated and deeply divergent lineages indicative of long-term persistence (Koskella et Brockhurst 2014; Trotereau et al. 2021). These differences were reflected in genome architecture. Lysogeny modules including integrases and excisionases were restricted to a limited subset of temperate phages (P68–P70), supporting site-specific chromosomal integration, whereas virulent lineages lacked canonical lysogeny modules but frequently encoded recombination-related genes such as transposases and genome rearrangement systems (Zhang et al. 2023; Pfeifer et Rocha 2024). The repeated detection of recombination-related genes in strictly lytic phages, coupled with the absence of integrases and excisionases, indicates that genome plasticity, rather than lysogeny, may constitute a primary mechanism of adaptive diversification in these lineages. This pattern suggests that RSSC phages can achieve evolutionary flexibility through ongoing genome rearrangements without adopting a temperate lifestyle. Functional repertoires followed the same dichotomy: virulent phages carried AMGs associated with rapid host exploitation and plant-associated substrates, whereas temperate phages encoded host interaction and persistence-related functions (Roux et al. 2016; Pfeifer et al. 2022). Together, these observations indicate that the RSSC phage community combines clonally expanding virulent lineages with persistent divergent phages, consistent with lifestyle-dependent evolutionary strategies structured by host ecology.

### A locally diversified and globally distinct phage assemblage

Genome comparisons showed that many Réunion Island phages lack close relatives among publicly available RSSC phages, with several forming monophyletic lineages consistent with novel taxa. This originality likely reflects the documented diversity of RSSC bacterial populations in the Southwest Indian Ocean (Prior et al. 2016; Yahiaoui et al. 2017; Cellier et al. 2023; Rasoamanana et al. 2020) combined with heterogeneous agroecological landscapes promoting diversification. Global genome relatedness analyses further supported this pattern: most Réunion Island phages formed coherent clusters separated from phages from other continents, including nearby Mauritius. Only a minority showed similarity to known lineages. The convergence of phylogenomic and global analyses indicates that the island constitutes a locally diversified evolutionary pool shaped by region-specific phage–bacteria interactions rather than passive sampling of widespread phages (Trotereau et al. 2021).

### Antibacterial efficacy depends on phage lifestyle and host phylogeny

*In vitro* inhibition was independent of plant origin but strongly associated with phage lifestyle. Strictly lytic phages showed consistently high antibacterial activity, whereas temperate phages produced weaker and more variable inhibition, consistent with previous reports of reduced biocontrol performance among lysogenic or temperate phages (Addy et al. 2012a,b; Álvarez et al. 2019). Genomic signatures supported these phenotypes: temperate phages encoded integrases related to site-specific chromosomal integration mechanisms (Panis et al. 2010), while virulent phages lacked complete lysogeny modules. In our study, virulent phages frequently encoded host exploitation functions such as pectin lyases, lipases, and DNA mimicry systems, whereas temperate phages encoded persistence-related genes including methyltransferases, involved in DNA methylation to protect the viral genome from host restriction systems, and nutrient acquisition functions. These differences provide a genomic context for reduced efficacy of temperate phages without implying direct causality. Host range assays further showed susceptibility structured by bacterial phylotype rather than geographic origin. Activity was restricted mainly to phylotype I and II-B.1 strains and absent against phylotypes III and IV, consistent with RSSC genetic differentiation (Prior et al. 2016). In Réunion Island, where phylotype I sequevar I-31 dominates (Yahiaoui et al. 2017; Cellier et al. 2023), the combination of narrow host range and strong inhibition among strictly lytic phages agrees with predictions that homogeneous host populations favor specialized, highly virulent phages (Torres-Barceló et al. 2025; Feltin et al. 2025). Together, infection success appears governed primarily by host evolutionary relatedness rather than spatial proximity, suggesting that effective biocontrol should prioritize adaptation to dominant bacterial lineages rather than broad host range.

### Evolution-informed selection and validation of a phage cocktail

Our five-phage cocktail combined strictly lytic phages with complementary host ranges and genomic repertoires. None encoded complete lysogeny modules, supporting their suitability for biocontrol (Holtappels et al. 2021). Several carried auxiliary functions previously associated with ecological performance in plant environments (Wang et al. 2019; Pfeifer et Rocha 2024). No antagonism was detected *in vitro*, and repeated applications provided stronger and more consistent protection against bacterial wilt *in planta* without phytotoxic effects, consistent with previous studies highlighting the importance of application strategy (Bae et al. 2012; Clavijo-Coppens et al. 2022). Beyond validating a specific formulation, these results demonstrate that no single experimental scale is sufficient to identify suitable biocontrol phages. *In vitro* assays revealed major differences in antibacterial efficacy and host range but required genomic data to distinguish virulent from temperate lifestyles and interpret functional repertoires. Conversely, genomic data alone was not predictive of phage performance, as related phages displayed contrasting inhibitory capacities. Effective selection therefore requires integrating complementary information: host range assays establish epidemiological relevance, virulence assays quantify antibacterial performance, and genome analyses ensure absence of lysogeny modules and toxin-releated systems, while providing ecological context (Hyman 2019). Combining phenotypic and genomic validation reduces the risk of selecting phages based solely on broad host range or novelty, which may not translate into biological efficacy (Torres-Barceló et al. 2025). This integrative framework supports rational cocktail design based on evolutionary compatibility and functional complementarity.

### Limitations, perspectives and conclusions

Our study presents several limitations. First, the use of a single bacterial strain for phage isolation may have biased recovery toward viruses infecting dominant host genotypes. Future work should therefore implement broader isolation strategies encompassing multiple strains and RSSC phylotypes to more accurately capture environmental diversity. Second, the evolution of bacterial resistance was not investigated, although it is a key determinant of long-term biocontrol efficacy (Labrie et al. 2010; Oechslin 2018). Subsequent studies should examine resistance trajectories, associated fitness trade-offs, and assess whether phage combinations can delay resistance emergence through complementary receptor targeting or coevolutionary dynamics. Prophage-mediated immunity may also modulate infection outcomes and should be considered in phage selection frameworks (Touchon et al. 2016). In addition, optimizing application parameters, including timing, frequency, and formulation will be critical to maximize field performance (Bae et al. 2012; Clavijo-Coppens et al. 2022). Expanding regional phage collections and incorporating rhizosphere microbial ecology will also further clarify mechanisms of disease suppression (Yang et al. 2023) and help define the spatial scale of local adaptation. Finally, extending comparable investigations to other crop pathogens beyond *Ralstonia*, such as soft-rot Pectobacteriaceae, will be particularly informative, as their contrasting infection strategies are likely to impose distinct selective pressures on associated phages, thereby refining our understanding of how pathogen ecology shapes phage diversity, specialization, and biocontrol potential.

In conclusion, our results indicate that Réunion Island hosts a locally diversified reservoir of RSSC phages primarily shaped by host population structure. The coexistence of expanding virulent lineages alongside persistent temperate phages suggests the presence of alternative evolutionary strategies maintained within a structured bacterial community. Antibacterial efficacy was strongly associated with evolutionary compatibility between phages and dominant host lineages, supporting the use of locally adapted phage cocktails. Altogether, these findings emphasize that effective and durable phage biocontrol relies on integrating ecological, evolutionary, and genomic insights rather than selecting candidates solely based on isolation source or taxonomic diversity.

## Data availability

All annotated Ralstonia phage genome sequences generated in this study were deposited in the European Nucleotide Archive (ENA) under BioProject accession number PRJEB111082 (see table S2 for the complete list of accession numbers and bioSample identifiers).

## Acknowledgments

We sincerely thank the Plant Protection Platform (3P, IBiSA), the members of producer cooperatives and farmers for their valuable assistance with sampling.

