## Supplementals for "Ecology, genomics and biocontrol potential of bacteriophages infecting the bacterial wilt pathogen *Ralstonia solanacearum* species complex in Réunion Island"

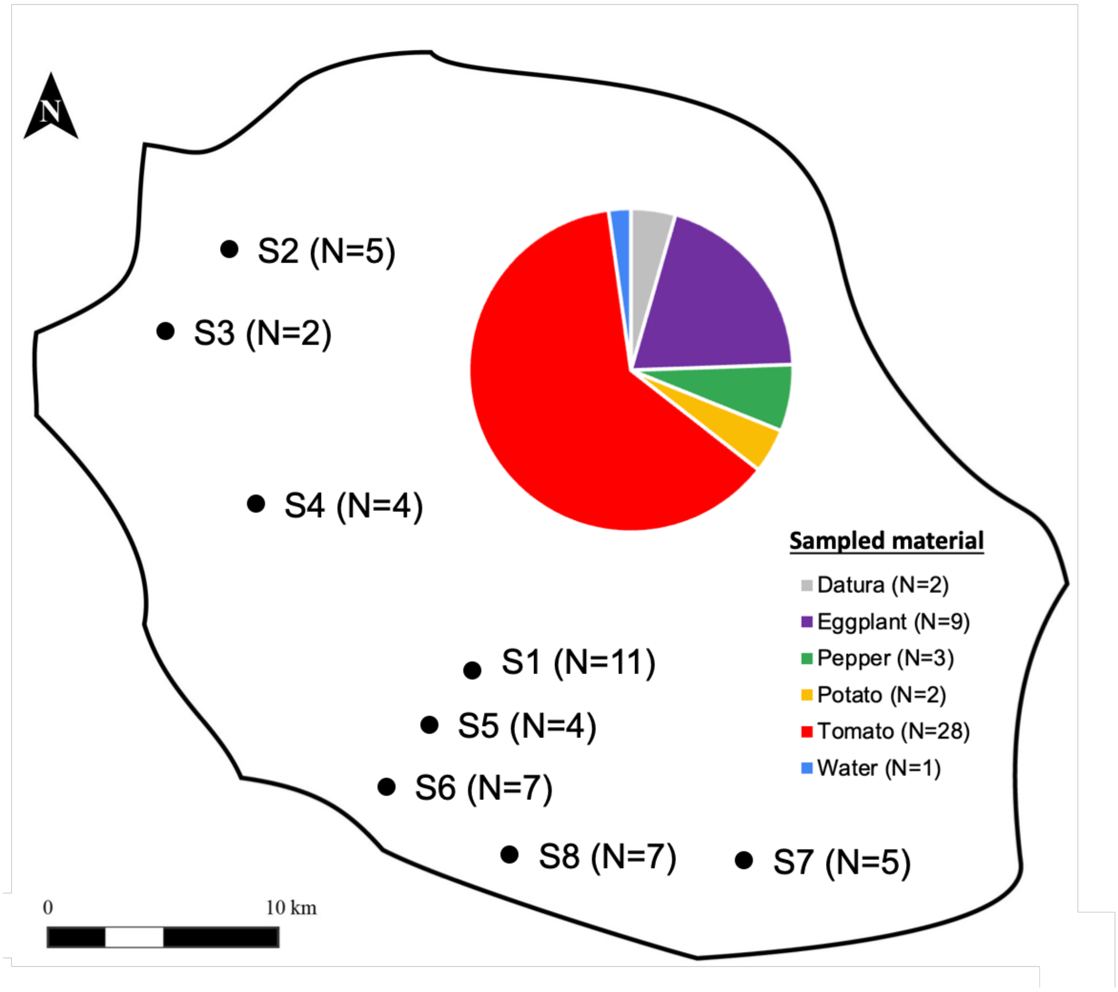


**FigureS1:** Geographic distribution of sampled sites and associated agricultural material.

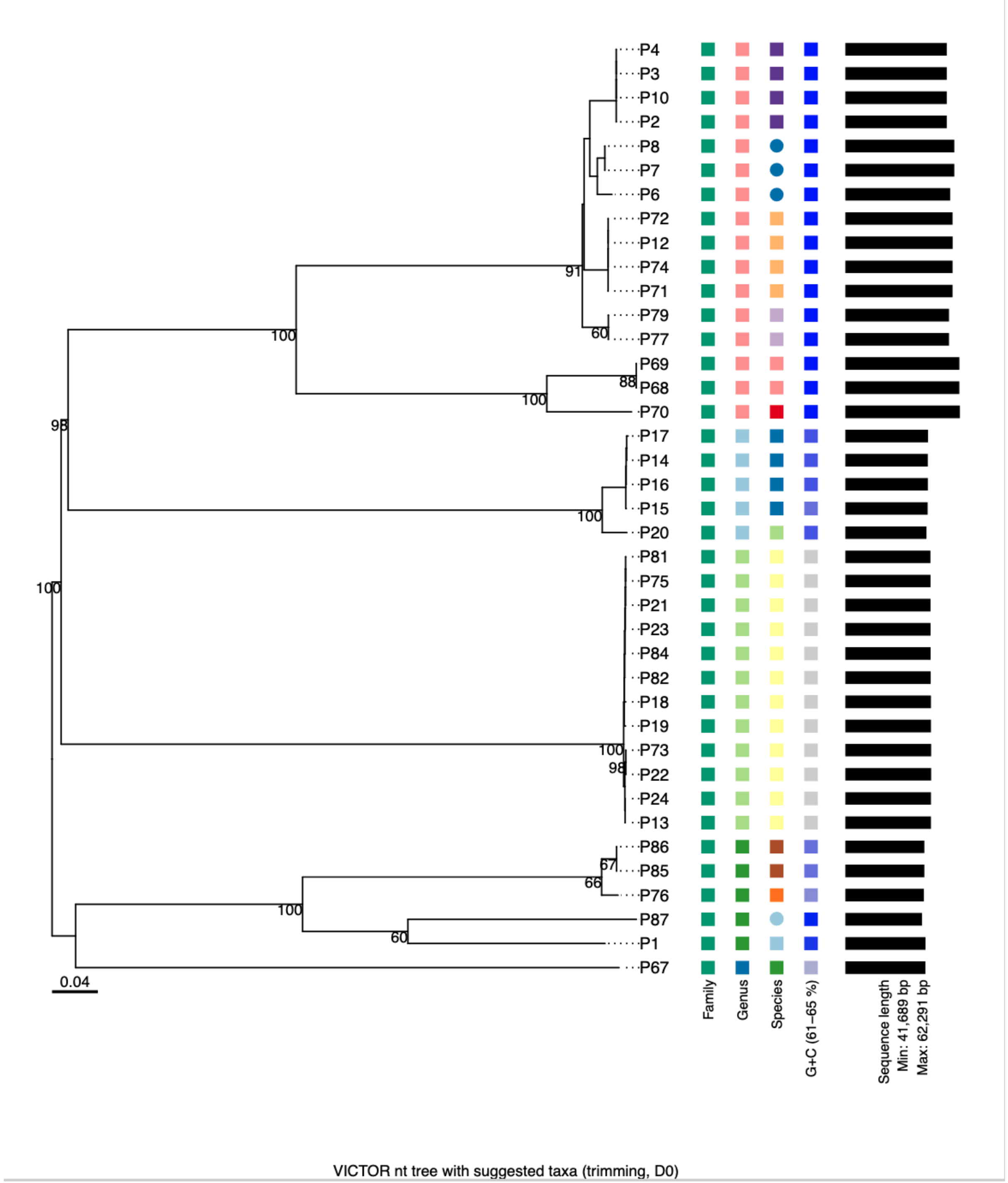


**FigureS2:** Phylogenetic tree based on intergenomic distances calculated with VICTOR (nucleotide-based, D0 formula, trimming enabled). Highly similar sequences were automatically removed.

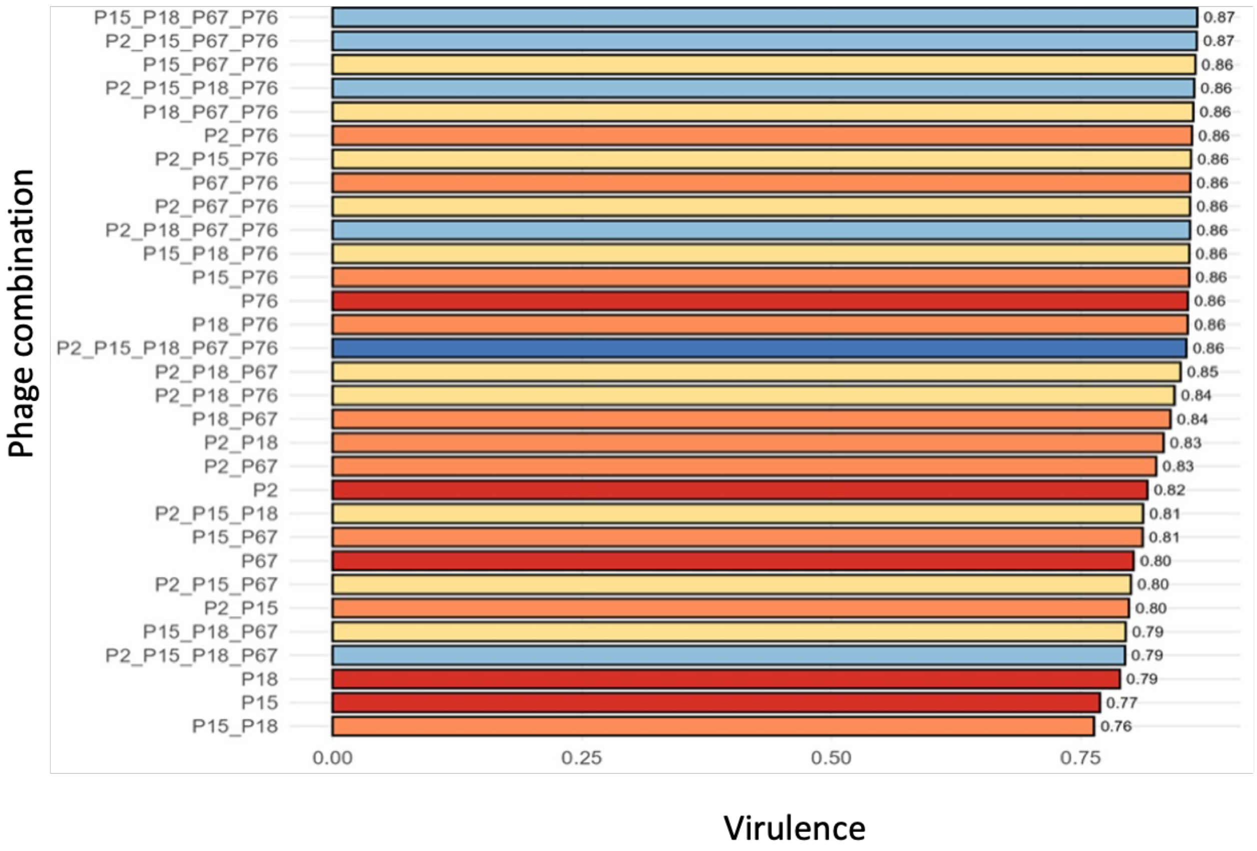


**FigureS3:** *In-vitro* assessment of the virulence of single phages and phage combinations against Ralstonia solanacearum

**Table S1**: Phylogenetic classification and geographic origin of the bacterial strains used in this study from the reference collection of the Pôle de Protection des Plantes (Réunion Island, France).

| Species | Strain | Phylotype | Sequevar | Year of isolation | Plant origin | Geographic origin | Refs |
|---|---|---|---|---|---|---|---|
| *Ralstonia solanacearum* | RUN0062 | IV | 10 | 1988 | Banana | Indonesia | (1). |
| *Ralstonia solanacearum* | RUN0320 | I | 46 | 2006 | Pepper | Madagascar | (1,2,3). |
| *Ralstonia solanacearum* | RUN0608 | I | 13 | 2000 | Potato | La Réunion | (1,2,4). |
| *Ralstonia solanacearum* | RUN2127 | I | 46 | 2012 | Tomato | Mayotte | (1,4,5) |
| *Ralstonia solanacearum* | RUN2280 | III | 59 | 2013 | Potato | Madagascar | (2). |
| *Ralstonia solanacearum* | RUN2340 | III | 58 | 2013 | Potato | Madagascar | (1,2). |
| *Ralstonia solanacearum* | RUN2510 | III | 60 | 2013 | Potato | Madagascar | (1,2). |
| *Ralstonia solanacearum* | RUN3012 | I | 31 | 2012 | Eggplant | La Réunion | (7). |
| *Ralstonia solanacearum* | RUN3149 | I | 18 | 2013 | Potato | Madagascar | (2,3,4) |
| *Ralstonia solanacearum* | RUN3570 | I | 33 | 2014 | Tomato | La Réunion | (7). |
| *Ralstonia solanacearum* | RUN3734 | III | 19 | 2014 | Tomato | La Réunion | (7). |
| *Ralstonia solanacearum* | RUN4269 | I | 14 | 2015 | Tomato | La Réunion | (7). |
| *Ralstonia solanacearum* | RUN4400 | I | 33 | 2015 | Tomato | Maurice | (1,4). |
| *Ralstonia solanacearum* | RUN4509 | IIB | 1 | 2015 | Potato | La Réunion | (1). |
| *Ralstonia solanacearum* | RUN4648 | I | 15 | 2015 | Potato | Maurice | (7). |
| *Ralstonia solanacearum* | RUN4847 | IV | 11 | 2015 | Potato | Maurice | (1). |
| *Ralstonia solanacearum* | RUN5350 | I | 64 | 2016 | Haricot | Maurice | (7). |
| *Ralstonia solanacearum* | RUN5405 | I | 18 | 2016 | Potato | Maurice | (7). |
| *Ralstonia solanacearum* | RUN5453 | I | 31 | 2016 | Tomato | Maurice | (1,4). |
| *Ralstonia solanacearum* | RUN6285 | I | 31 | 2019 | Eggplant | Madagascar | (3). |
| *Ralstonia solanacearum* | RUN6499 | I | 18 | 2019 | Eggplant | Madagascar | (3). |
| *Ralstonia solanacearum* | RUN6514 | IIB | 1 | 2019 | Potato | Madagascar | (7). |
| *Ralstonia solanacearum* | RUN6534 | I | 18 | 2019 | Pepper | Madagascar | (3). |
| *Ralstonia solanacearum* | RUN6565 | III | 19 | 2019 | Haricot | Madagascar | (7). |
| *Ralstonia solanacearum* | RUN2161 | I | 18 | 2012 | Tomato | Mayotte | (5,6). |

(3). *doi.org/10.1094/PHYTOFR-10-24-0110-A*

(4). *doi.org/10.1371/journal.pone.0242846*

(5). *10.3389/fpls.2017.02209*

(6). *10.1186/s12863-025-01395-2*

(7). Unpublished data

**Table S2.** Accession numbers and BioSample identifiers of the samples and genomes generated in this study. New accession numbers are provided for 10 genomes previously published, following re-annotation as described in the main text.

| Isolate | Previous genome accession number | New genome acession number | New bioSample number |
|---|---|---|---|
| P1 | MT740728 | ERS30300663 | SAMEA122677856 |
| P2 | MT740730 | ERS30300668 | SAMEA122677861 |
| P3 | **This study** | ERS30300669 | SAMEA122677862 |
| P4 | **This study** | ERS30300670 | SAMEA122677863 |
| P5 | **This study** | ERS30300671 | SAMEA122677864 |
| P6 | MT740733 | ERS30300672 | SAMEA122677865 |
| P7 | **This study** | ERS30300673 | SAMEA122677866 |
| P8 | **This study** | ERS30300674 | SAMEA122677867 |
| P9 | **This study** | ERS30300675 | SAMEA122677868 |
| P10 | **This study** | ERS30300676 | SAMEA122677869 |
| P11 | **This study** | ERS30300677 | SAMEA122677870 |
| P12 | MT740742 | ERS30300678 | SAMEA122677871 |
| P13 | **This study** | ERS30300687 | SAMEA122677880 |
| P14 | MT740747 | ERS30300664 | SAMEA122677857 |
| P15 | MT740725 | ERS30300665 | SAMEA122677858 |
| P16 | MT740746 | ERS30300666 | SAMEA122677859 |
| P17 | MT740735 | ERS30300667 | SAMEA122677860 |
| P18 | **This study** | ERS30300688 | SAMEA122677881 |
| P19 | **This study** | ERS30300689 | SAMEA122677882 |
| P20 | **This study** | ERS30302057 | SAMEA122679250 |
| P21 | MT740745 | ERS30300690 | SAMEA122677883 |
| P22 | **This study** | ERS30300691 | SAMEA122677884 |
| P23 | **This study** | ERS30300692 | SAMEA122677885 |
| P24 | MT740726 | ERS30300693 | SAMEA122677886 |
| P67 | **This study** | ERS30300686 | SAMEA122677879 |
| P68 | **This study** | ERS30302058 | SAMEA122679251 |
| P69 | **This study** | ERS30302059 | SAMEA122679252 |
| P70 | **This study** | ERS30302060 | SAMEA122679253 |
| P71 | **This study** | ERS30300679 | SAMEA122677872 |
| P72 | **This study** | ERS30300680 | SAMEA122677873 |
| P73 | **This study** | ERS30300694 | SAMEA122677887 |
| P74 | **This study** | ERS30300681 | SAMEA122677874 |
| P75 | **This study** | ERS30300695 | SAMEA122677888 |
| P76 | **This study** | ERS30302061 | SAMEA122679254 |
| P77 | **This study** | ERS30300682 | SAMEA122677875 |
| P78 | **This study** | ERS30300683 | SAMEA122677876 |
| P79 | **This study** | ERS30300684 | SAMEA122677877 |
| P80 | **This study** | ERS30300685 | SAMEA122677878 |
| P81 | **This study** | ERS30300696 | SAMEA122677889 |
| P82 | **This study** | ERS30300697 | SAMEA122677890 |
| P83 | **This study** | ERS30300698 | SAMEA122677891 |
| P84 | **This study** | ERS30300699 | SAMEA122677892 |
| P85 | **This study** | ERS30302055 | SAMEA122679248 |
| P86 | **This study** | ERS30302056 | SAMEA122679249 |
| P87 | **This study** | ERS30302062 | SAMEA122679255 |

**Table S3.** Curated dataset of complete Caudovirales phage genomes infecting members of the order Burkholderiales available in GenBank (1/2)

| N | Organism_Name | Country | Host | Species | Genus | Family | Accession | Assembly_Accession |
|---|---|---|---|---|---|---|---|---|
| 1 | Burkholderia phage AMP1 | NA | Burkholderia pseudomallei | Ampunavirus BpAMP1 | Ampunavirus | Autographiviridae | GCA_008605765.1 | ASM860576v1 |
| 2 | Burkholderia phage AP3 | NA | Burkholderia cenocepacia | Aptresvirus AP3 | Aptresvirus | Peduoviridae | NC_047752 | GCA_002605605.1 |
| 3 | Burkholderia phage BCSR129 | Israel | Burkholderia | Burkholderia phage BCSR129 | NA | NA | MW460247 | GCA_016865795.1 |
| 4 | Burkholderia phage BCSR5 | Israel | Burkholderia | Burkholderia phage BCSR5 | NA | NA | MW460245 | GCA_016865805.1 |
| 5 | Burkholderia phage BCSR52 | Israel | Burkholderia | Burkholderia phage BCSR52 | Bcepfunavirus | NA | GCA_016865815.1 | ASM1686581v1 |
| 6 | Burkholderia phage BEK | Pakistan | Burkholderia pseudomallei 9 | Tigrvirus phi52237 | Tigrvirus | Peduoviridae | GCA_002756315.1 | ASM275631v1 |
| 7 | Burkholderia phage Bcep1 | NA | Burkholderia cenocepacia | Naesvirus bcep1 | Naesvirus | NA | NC_005263 | GCA_000845445.1 |
| 8 | Burkholderia phage Bcep176 | NA | Burkholderia cepacia | Burkholderia phage Bcep176 | Stanholtvirus | NA | GCA_000865925.1 | ViralProj16102 |
| 9 | Burkholderia phage Bcep22 | NA | Burkholderia | Lessievirus bcep22 | Lessievirus | NA | NC_005262 | GCA_000840865.2 |
| 10 | Burkholderia phage Bcep43 | NA | Burkholderia cepacia | Naesvirus bcep43 | Naesvirus | NA | NC_005342 | GCA_000844585.1 |
| 11 | Burkholderia phage Bcep781 | NA | Burkholderia cepacia | Naesvirus bcep781 | Naesvirus | NA | NC_004333 | GCA_000845425.1 |
| 12 | Burkholderia phage BcepB1A | NA | Burkholderia | Burkholderia phage BcepB1A | NA | NA | NC_005886 | GCA_000844905.1 |
| 13 | Burkholderia phage BcepC6B | NA | Burkholderia cepacia | Ryyoungvirus bcepC6B | Ryyoungvirus | NA | GCA_000843605.1 | ViralProj14379 |
| 14 | Burkholderia phage BcepF1 | NA | Burkholderia ambifaria | Bcepfunavirus bcepF1 | Bcepfunavirus | NA | GCA_000873005.1 | ViralProj18857 |
| 15 | Burkholderia phage BcepGomr | NA | Burkholderia cepacia | Burkholderia phage BcepGomr | NA | NA | NC_009447 | GCA_000873805.1 |
| 16 | Burkholderia phage BcepIL02 | NA | Burkholderia cenocepacia | Lessievirus bcepil02 | Lessievirus | NA | NC_012743 | GCA_000882995.1 |
| 17 | Burkholderia phage BcepMigl | NA | Burkholderia cenocepacia | Lessievirus bcepmigl | Lessievirus | NA | NC_019917 | GCA_000903775.1 |
| 18 | Burkholderia phage BcepMu | NA | Burkholderia | Bcepmuvirus bcepMu | Bcepmuvirus | 1 | GCA_000841805.1 | ViralProj14376 |
| 19 | Burkholderia phage BcepNY3 | NA | Burkholderia cenocepacia HI2424 | Naesvirus bcepNY3 | Naesvirus | NA | NC_009604 | GCA_000873385.1 |
| 20 | Burkholderia phage BcepNazgul | NA | Burkholderia cepacia | Burkholderia virus BcepNazgul | Nazgulvirus | Casjensviridae | GCA_000840725.1 | ViralProj14305 |
| 21 | Burkholderia phage BcepSaruman | USA | Burkholderia cenocepacia | Sarumanvirus bcepsaruman | Sarumanvirus | NA | GCA_004771375.1 | ASM477137v1 |
| 22 | Burkholderia phage BcepSauron | USA | Burkholderia cenocepacia | Sarumanvirus bcepsauron | Sarumanvirus | NA | GCA_005394215.1 | ASM539421v1 |
| 23 | Burkholderia phage BgManors32 | Israel | Burkholderia gladioli | Bigmanorsvirus bgmanors32 | Bigmanorsvirus | NA | GCA_021355205.1 | ASM2135520v1 |
| 24 | Burkholderia phage BgVeeders33 | Israel | Burkholderia gladioli | Burkholderia phage BgVeeders33 | NA | NA | OK665841 | GCA_021355195.1 |
| 25 | Burkholderia phage Bm1 | NA | Burkholderia multivorans | Burkholderia phage Bm1 | NA | NA | PP183292 | GCA_036852335.1 |
| 26 | Burkholderia phage Bp-AMP1 | Thailand | Burkholderia pseudomallei K96243 | Ampunavirus BpAMP1 | Ampunavirus | Autographiviridae | GCA_002604225.1 | ASM260422v1 |
| 27 | Burkholderia phage Bp-AMP2 | Thailand | Burkholderia pseudomallei K96243 | Ampunavirus BpAMP1 | Ampunavirus | Autographiviridae | GCA_002604265.1 | ASM260426v1 |
| 28 | Burkholderia phage Bp-AMP3 | Thailand | Burkholderia pseudomallei K96243 | Ampunavirus BpAMP1 | Ampunavirus | Autographiviridae | GCA_002606825.1 | ASM260682v1 |
| 29 | Burkholderia phage Bp-AMP4 | Thailand | Burkholderia pseudomallei K96243 | Ampunavirus BpAMP1 | Ampunavirus | Autographiviridae | GCA_002604285.1 | ASM260428v1 |
| 30 | Burkholderia phage Bups phi1 | Australia | Burkholderia pseudomallei NCTC 13178 | Burkholderia phage Bups phi1 | NA | NA | EU307291 | GCA_009673985.1 |
| 31 | Burkholderia phage CSP3 | Australia | Burkholderia contaminans | Burkholderia phage CSP3 | Lessievirus | NA | OQ053201 | GCA_027618055.1 |
| 32 | Burkholderia phage Carl1 | Canada | Burkholderia cenocepacia | Burkholderia phage Carl1 | Kisquinquevirus | Peduoviridae | GCA_024023325.1 | ASM2402332v1 |
| 33 | Burkholderia phage DC1 | NA | Burkholderia cenocepacia | Lessievirus DC1 | Lessievirus | NA | NC_018452 | GCA_000899095.1 |
| 34 | Burkholderia phage FLC10 | Japan | Burkholderia glumae | Kisquattuordecimvirus FLC10 | Kisquattuordecimvirus | Peduoviridae | GCA_021355115.1 | ASM2135511v1 |
| 35 | Burkholderia phage FLC5 | Japan | Burkholderia glumae | Kisquattuordecimvirus FLC5 | Kisquattuordecimvirus | Peduoviridae | GCA_014488745.1 | ASM1448874v1 |
| 36 | Burkholderia phage FLC6 | NA | Burkholderia glumae | Burkholderia phage FLC6 | Chiangmaivirus | NA | GCA_016811315.1 | ASM1681131v1 |
| 37 | Burkholderia phage FLC8 | Japan | Burkholderia glumae | Burkholderia phage FLC8 | Chiangmaivirus | NA | GCA_021355095.1 | ASM2135509v1 |
| 38 | Burkholderia phage FLC9 | Japan | Burkholderia glumae | Burkholderia phage FLC9 | NA | NA | LC667451 | GCA_021355105.1 |
| 39 | Burkholderia phage JC1 | Canada | Burkholderia cenocepacia | Burkholderia phage JC1 | Lessievirus | NA | OM283127 | GCA_022516615.1 |
| 40 | Burkholderia phage JG068 | NA | Burkholderia cenocepacia | Mguuvirus JG068 | Mguuvirus | Autographiviridae | NC_022916 | GCA_000914995.1 |
| 41 | Burkholderia phage KL3 | NA | Burkholderia cenocepacia | Kayeltresvirus KL3 | Kayeltresvirus | Peduoviridae | NC_015266 | GCA_000893295.1 |
| 42 | Burkholderia phage KS10 | NA | Burkholderia cenocepacia | Burkholderia phage KS10 | NA | NA | NC_011216 | GCA_000881475.1 |
| 43 | Burkholderia phage KS14 | NA | Burkholderia ambifaria | Kisquattuordecimvirus KS14 | Kisquattuordecimvirus | Peduoviridae | NC_015273 | GCA_000891775.1 |
| 44 | Burkholderia phage KS5 | NA | Burkholderia cenocepacia | Kisquinquevirus KS5 | Kisquinquevirus | Peduoviridae | NC_015265 | GCA_000891735.1 |
| 45 | Burkholderia phage KS9 | Canada | Burkholderia pseudomallei | Burkholderia phage KS9 | Stanholtvirus | 1 | GCA_000885155.1 | ViralProj39771 |
| 46 | Burkholderia phage Magia | USA | Burkholderia cenocepacia | Magiavirus magia | Magiavirus | NA | NC_070954 | GCA_015502015.1 |
| 47 | Burkholderia phage Maja | USA | Burkholderia gladioli | Burkholderia phage Maja | Bcepfunavirus | NA | GCA_015244865.1 | ASM1524486v1 |
| 48 | Burkholderia phage Mana | USA | Burkholderia gladioli | Aptresvirus mana | Aptresvirus | Peduoviridae | GCA_015502025.1 | ASM1550202v1 |
| 49 | Burkholderia phage Menos | USA | Burkholderia cenocepacia | Burkholderia phage Menos | NA | Peduoviridae | GCA_022818335.1 | ASM2281833v1 |
| 50 | Burkholderia phage Mica | USA | Burkholderia cenocepacia | Micavirus Mica | Micavirus | NA | NC_054148 | GCA_015502035.1 |
| 51 | Burkholderia phage Milagro | USA | Burkholderia cenocepacia | Burkholderia phage Milagro | NA | Peduoviridae | GCA_022818345.1 | ASM2281834v1 |
| 52 | Burkholderia phage Momento | USA | Burkholderia cenocepacia | Burkholderia phage Momento | NA | Peduoviridae | GCA_022818355.1 | ASM2281835v1 |
| 53 | Burkholderia phage Musica | USA | Burkholderia cenocepacia | Burkholderia phage Musica | NA | Peduoviridae | GCA_022818365.1 | ASM2281836v1 |
| 54 | Burkholderia phage PE067 | Germany | Burkholderia thailandensis | Burkholderia phage PE067 | NA | NA | KT803877 | GCA_009901735.1 |
| 55 | Burkholderia phage PK23 | Thailand | Burkholderia pseudomallei | Burkholderia phage PK23 | Duodecimduovirus | Peduoviridae | GCA_019095875.1 | ASM1909587v1 |
| 56 | Burkholderia phage Paku | USA | Burkholderia cenocepacia | Burkholderia phage Paku | NA | Autographiviridae | GCA_020484125.1 | ASM2048412v1 |
| 57 | Burkholderia phage PhiBP82.1 | NA | Burkholderia pseudomallei | Stanholtvirus sv1026b | Stanholtvirus | NA | GCA_022694595.1 | ASM2269459v1 |
| 58 | Burkholderia phage PhiBP82.2 | NA | Burkholderia pseudomallei | Tigrvirus BP822 | Tigrvirus | Peduoviridae | GCA_022213645.1 | ASM2221364v1 |
| 59 | Burkholderia phage PhiBP82.3 | NA | Burkholderia pseudomallei | Burkholderia phage PhiBP82.3 | Tigrvirus | Peduoviridae | GCA_023170455.1 | ASM2317045v1 |
| 60 | Burkholderia phage PhiBt-E264.1 | NA | Burkholderia pseudomallei | Burkholderia phage PhiBt-E264.1 | Stanholtvirus | NA | GCA_025787965.1 | ASM2578796v1 |
| 61 | Burkholderia phage ST79 | NA | Burkholderia pseudomallei | Nampongvirus ST79 | Nampongvirus | Peduoviridae | GCA_000908615.1 | ViralProj206488 |
| 62 | Burkholderia phage phi1026b | NA | Burkholderia pseudomallei 1026b | Stanholtvirus sv1026b | Stanholtvirus | 1 | GCA_000846305.1 | ViralProj14410 |
| 63 | Burkholderia phage phi644-2 | NA | Burkholderia pseudomallei | Stanholtvirus sv6442 | Stanholtvirus | NA | GCA_000893155.1 | ViralProj62941 |
| 64 | Burkholderia phage phiBt-TXDOH | USA | Burkholderia pseudomallei | Burkholderia phage phiBt-TXDOH | Stanholtvirus | NA | GCA_020523175.1 | ASM2052317v1 |
| 65 | Burkholderia phage phiBtTUL1a | NA | Burkholderia pseudomallei | Burkholderia phage phiBtTUL1a | Stanholtvirus | NA | GCA_032446275.1 | ASM3244627v1 |
| 66 | Burkholderia phage phiE058 | Germany | Burkholderia thailandensis | Burkholderia phage phiE058 | NA | NA | MH809533 | GCA_003668395.1 |
| 67 | Burkholderia phage phiE094 | Thailand | Burkholderia thailandensis | Tigrvirus phiE094 | Tigrvirus | Peduoviridae | GCA_017654305.1 | ASM1765430v1 |
| 68 | Burkholderia phage phiE12-2 | NA | Burkholderia pseudomallei | Duodecimduovirus phiE122 | Duodecimduovirus | Peduoviridae | GCA_000871505.1 | ViralProj19161 |
| 69 | Burkholderia phage phiE125 | NA | Burkholderia thailandensis | Stanholtvirus E125 | Stanholtvirus | 1 | GCA_000840845.1 | ViralProj14330 |
| 70 | Burkholderia phage phiE131 | Germany | Burkholderia thailandensis | Burkholderia phage phiE131 | NA | NA | MH809532 | GCA_003668375.1 |
| 71 | Burkholderia phage phiE202 | NA | Burkholderia thailandensis | Tigrvirus E202 | Tigrvirus | Peduoviridae | GCA_000870585.1 | ViralProj19163 |
| 72 | Burkholderia phage phiE255 | NA | Burkholderia thailandensis | Bcepmuvirus E255 | Bcepmuvirus | NA | GCA_000873105.1 | ViralProj19165 |
| 73 | Burkholderia phage phiE52237 | NA | Burkholderia pseudomallei | Tigrvirus phi52237 | Tigrvirus | Peduoviridae | GCA_000861045.1 | ViralProj15422 |
| 74 | Burkholderia phage phiX216 | Thailand | Burkholderia cepacia | Tigrvirus phi52237 | Tigrvirus | Peduoviridae | JX681814 | GCA_002755455.1 |
| 75 | Burkholderia phage vB_BceS_AH2 | NA | Burkholderia ambifaria | Burkholderia virus AH2 | Ahduovirus | Casjensviridae | NC_018283 | GCA_000898995.1 |
| 76 | Burkholderia phage vB_BceS_KL1 | NA | Burkholderia ambifaria | Kilunavirus KL1 | Kilunavirus | NA | NC_018278 | GCA_000897395.1 |
| 77 | Burkholderia phage vB_BmuP_KL4 | NA | Burkholderia | Kelquatrovirus KL4 | Kelquatrovirus | NA | GCA_003094055.1 | ASM309405v1 |
| 78 | Burkholderia phage vB_BpP_HN01 | China | Burkholderia pseudomallei | Burkholderia phage vB_BpP_HN01 | NA | Schitoviridae | GCA_023239875.1 | ASM2323987v1 |
| 79 | Burkholderia phage vB_BpP_HN02 | China | Burkholderia | Burkholderia phage vB_BpP_HN02 | NA | Schitoviridae | GCA_036429425.1 | ASM3642942v1 |

**Table S3.** Curated dataset of complete Caudovirales phage genomes infecting members of the order Burkholderiales available in GenBank (2/2)

| N | Organism_Name | Country | Host | Species | Genus | Family | Accession | Assembly_Accession |
|---|---|---|---|---|---|---|---|---|
| 80 | Burkholderia phage vB_BpP_HN03 | China | Burkholderia pseudomallei | Burkholderia phage vB_BpP_HN03 | NA | NA | PP100843 | GCA_036924145.1 |
| 81 | Burkholderia phage vB_BpP_HN04 | China | Burkholderia pseudomallei | Burkholderia phage vB_BpP_HN04 | NA | NA | PP100844 | GCA_036924155.1 |
| 82 | Burkholderia phage vB_BpP_HN05 | China | Burkholderia pseudomallei | Burkholderia phage vB_BpP_HN05 | NA | NA | PP100845 | GCA_036924165.1 |
| 83 | Burkholderia phage vB_HM387 | Thailand | Burkholderia pseudomallei | Burkholderia phage vB_HM387 | Tigrvirus | Peduoviridae | GCA_038023155.1 | ASM3802315v1 |
| 84 | Burkholderia phage vB_HM795 | Thailand | Burkholderia pseudomallei | Burkholderia phage vB_HM795 | NA | NA | OR990505 | GCA_038023165.1 |
| 85 | Burkholderiales phage 68_11 | USA | Burkholderiales | Burkholderiales phage 68_11 | NA | NA | MN304815 | GCA_020473115.1 |
| 86 | Ralstonia phage 10RS305A | China | Ralstonia solanacearum | Ralstonia phage 10RS305A | Serkorvirus | Autographiviridae | GCA_025790075.1 | ASM2579007v1 |
| 87 | Ralstonia phage 10RS306A | China | Ralstonia solanacearum | Ralstonia phage 10RS306A | Serkorvirus | Autographiviridae | GCA_025630445.1 | ASM2563044v1 |
| 88 | Ralstonia phage Adzire | Reunion | Ralstonia pseudosolanacearum | Bakolyvirus simangalove | Bakolyvirus | NA | NC_054462 | GCA_014262625.1 |
| 89 | Ralstonia phage AhaGv | China | Ralstonia solanacearum | Ralstonia phage AhaGv | Gervaisevirus | NA | GCA_035764955.1 | ASM3576495v1 |
| 90 | Ralstonia phage Albius | Reunion | Ralstonia pseudosolanacearum | Rahariannevirus raharianne | Rahariannevirus | NA | GCA_014262695.1 | ASM1426269v1 |
| 91 | Ralstonia phage Alix | Mauritius | Ralstonia pseudosolanacearum | Gervaisevirus claudettte | Gervaisevirus | NA | GCA_014262865.1 | ASM1426286v1 |
| 92 | Ralstonia phage Anchaing | Reunion | Ralstonia pseudosolanacearum | Anchaingvirus anchaing | Anchaingvirus | Autographiviridae | GCA_014262885.1 | ASM1426288v1 |
| 93 | Ralstonia phage Bakoly | Mauritius | Ralstonia pseudosolanacearum | Bakolyvirus bakoly | Bakolyvirus | NA | NC_054463 | GCA_014262575.1 |
| 94 | Ralstonia phage BHDT_So9 | Viet Nam | Ralstonia solanacearum | Ralstonia phage BHDT_So9 | Higashivirus | Autographiviridae | GCA_025204285.1 | ASM2520428v1 |
| 95 | Ralstonia phage BHDT8 | Viet Nam | Ralstonia solanacearum | Ralstonia phage BHDT8 | Higashivirus | Autographiviridae | GCA_029376405.1 | ASM2937640v1 |
| 96 | Ralstonia phage BHDTSo81 | Viet Nam | Ralstonia solanacearum | Ralstonia phage BHDTSo81 | Higashivirus | Autographiviridae | GCA_026432335.1 | ASM2643233v1 |
| 97 | Ralstonia phage BOESR1 | Thailand | Ralstonia pseudosolanacearum | Ralstonia phage BOESR1 | Gyeongsanvirus | Autographiviridae | ASM2937643v1 | NA |
| 98 | Ralstonia phage Cimandef | Reunion | Ralstonia pseudosolanacearum | Cimandefvirus cimandef | Cimandefvirus | NA | GCA_014262965.2 | ASM1426296v2 |
| 99 | Ralstonia phage Claudette | Mauritius | Ralstonia pseudosolanacearum | Gervaisevirus claudettte | Gervaisevirus | NA | GCA_014263015.2 | ASM1426301v2 |
| 100 | Ralstonia phage Darius | Mauritius | Ralstonia pseudosolanacearum | Gervaisevirus gervaise | Gervaisevirus | NA | GCA_014263035.1 | ASM1426303v1 |
| 101 | Ralstonia phage Dimitile | Mauritius | Ralstonia pseudosolanacearum | Ralstonia phage Dimitile | Cimandefvirus | NA | GCA_014263055.1 | ASM1426305v1 |
| 102 | Ralstonia phage Dina | Mauritius | Ralstonia pseudosolanacearum | Dinavirus dina | Dinavirus | NA | NC_055026 | GCA_014263305.1 |
| 103 | Ralstonia phage DLDT_So2 | Viet Nam | Ralstonia solanacearum | Ralstonia phage DLDT_So2 | Higashivirus | Autographiviridae | GCA_025204295.1 | ASM2520429v1 |
| 104 | Ralstonia phage DU_RP_I | South Korea | Ralstonia solanacearum | Gyeongsanvirus DURPI | Gyeongsanvirus | Autographiviridae | GCA_002956085.1 | ASM295608v1 |
| 105 | Ralstonia phage DU_RP_II | NA | Ralstonia solanacearum | Ralstonia phage DU_RP_II | NA | NA | GCA_002627225.1 | ASM262722v1 |
| 106 | Ralstonia phage Elie | Reunion | Ralstonia pseudosolanacearum | Ralstonia phage Elie | Bakolyvirus | NA | MT740735 | GCA_014262725.1 |
| 107 | Ralstonia phage Eline | Mauritius | Ralstonia pseudosolanacearum | Cimandefvirus eline | Cimandefvirus | NA | GCA_014263095.1 | ASM1426309v1 |
| 108 | Ralstonia phage Firinga | Mauritius | Ralstonia pseudosolanacearum | Firingavirus firinga | Firingavirus | NA | NC_054961 | GCA_014263175.1 |
| 109 | Ralstonia phage Gamede | Mauritius | Ralstonia pseudosolanacearum | Cimandefvirus gamede | Cimandefvirus | NA | GCA_014263205.2 | ASM1426320v2 |
| 110 | Ralstonia phage Gerry | Mauritius | Ralstonia pseudosolanacearum | Cimandefvirus Ggerry | Cimandefvirus | NA | GCA_014263215.1 | ASM1426321v1 |
| 111 | Ralstonia phage Gervaise | Mauritius | Ralstonia pseudosolanacearum | Gervaisevirus gervaise | Gervaisevirus | NA | GCA_014263265.1 | ASM1426326v1 |
| 112 | Ralstonia phage GP4 | NA | Ralstonia solanacearum | Gervaisevirus GP4 | Gervaisevirus | NA | GCA_003354205.1 | ASM335420v1 |
| 113 | Ralstonia phage Hennie | Mauritius | Ralstonia pseudosolanacearum | Firingavirus firinga | Firingavirus | NA | MT740741 | GCA_014263275.1 |
| 114 | Ralstonia phage Heva | Reunion | Ralstonia pseudosolanacearum | Cimandefvirus heva | Cimandefvirus | NA | GCA_014263285.1 | ASM1426328v1 |
| 115 | Ralstonia phage Hyacinthe | Mauritius | Ralstonia pseudosolanacearum | Rahariannevirus raharianne | Rahariannevirus | NA | GCA_014263295.1 | ASM1426329v1 |
| 116 | Ralstonia phage Jenny | Mauritius | Ralstonia pseudosolanacearum | Bakolyvirus bakoly | Bakolyvirus | NA | MT740744 | GCA_014262735.1 |
| 117 | Ralstonia phage p2106 | China | Ralstonia solanacearum | Ralstonia phage p2106 | Serkorvirus | Autographiviridae | GCA_027574145.2 | ASM2757414v1 |
| 118 | Ralstonia phage p2110 | China | Ralstonia solanacearum | Ralstonia phage p2110 | Gervaisevirus | NA | GCA_027574155.2 | ASM2757415v1 |
| 119 | Ralstonia phage p2137 | China | Ralstonia solanacearum | Ralstonia phage p2137 | Serkorvirus | Autographiviridae | GCA_027574165.2 | ASM2757416v1 |
| 120 | Ralstonia phage phiAp1 | Brazil | Ralstonia solanacearum | Ayakvirus Ap1 | Ayakvirus | Autographiviridae | GCA_002617905.1 | ASM261790v1 |
| 121 | Ralstonia phage phiITL-1 | Mexico | Ralstonia solanacearum | Serkorvirus ITL1 | Serkorvirus | Autographiviridae | GCA_002605405.1 | ITL-1 |
| 122 | Ralstonia phage phiRSL1 | NA | Ralstonia | Mieseafarmvirus RSL1 | Mieseafarmvirus | 1 | GCA_000879455.1 | ViralProj30059 |
| 123 | Ralstonia phage phiRSP | Mexico | Ralstonia solanacearum | Coatlandelriovirus RSP | Coatlandelriovirus | NA | GCA_003368625.1 | ASM336862v1 |
| 124 | Ralstonia phage P-PSG-11 | China | Ralstonia solanacearum | Ralstonia phage P-PSG-11 | Gyeongsanvirus | Autographiviridae | GCA_009185825.1 | ASM918582v1 |
| 125 | Ralstonia phage P-PSG-11-1 | China | Ralstonia solanacearum | Ralstonia phage P-PSG-11-1 | Gyeongsanvirus | Autographiviridae | GCA_009185865.1 | ASM918586v1 |
| 126 | Ralstonia phage PQ43W | China | Ralstonia pseudosolanacearum | Ralstonia phage PQ43W | NA | NA | PP405626 | GCA_037061965.1 |
| 127 | Ralstonia phage QkW1 | China | Ralstonia pseudosolanacearum | Ralstonia phage QkW1 | NA | NA | PP236382 | GCA_036630285.1 |
| 128 | Ralstonia phage Raharianne | Reunion | Ralstonia pseudosolanacearum | Rahariannevirus raharianne | Rahariannevirus | NA | GCA_014262805.1 | ASM1426280v1 |
| 129 | Ralstonia phage Reminis | NA | Ralstonia | Reminisvirus reminis | Reminisvirus | Autographiviridae | MT331608 | GCA_009667685.1 |
| 130 | Ralstonia phage RP12 | Thailand | Ralstonia solanacearum | Ripduovirus RP12 | Ripduovirus | NA | NC_041911 | GCA_002617345.1 |
| 131 | Ralstonia phage RP13 | NA | Ralstonia solanacearum | Ralstonia phage RP13 | NA | NA | LC554890 | GCA_013340905.1 |
| 132 | Ralstonia phage RP31 | Thailand | Ralstonia solanacearum | Ripduovirus RP12 | Ripduovirus | NA | AP017925 | GCA_002617365.1 |
| 133 | Ralstonia phage RPSC1 | China | Ralstonia solanacearum | Stompelvirus RPSC1 | Stompelvirus | Autographiviridae | NC_047982 | GCA_003288515.1 |
| 134 | Ralstonia phage RpT1 | South Korea | Ralstonia pseudosolanacearum | Ralstonia phage RpT1 | Serkorvirus | Autographiviridae | OK274245 | GCA_020523225.1 |
| 135 | Ralstonia phage RpY1 | NA | Ralstonia solanacearum | Ralstonia phage RpY1 | NA | NA | MN996301 | GCA_014071215.1 |
| 136 | Ralstonia phage RpY2 | South Korea | Ralstonia pseudosolanacearum | Ralstonia phage RpY2 | Serkorvirus | Autographiviridae | OK318991 | GCA_020523285.1 |
| 137 | Ralstonia phage RPZH3 | China | Ralstonia solanacearum | Ralstonia phage RPZH3 | Gervaisevirus | NA | GCA_026568075.1 | ASM2656807v1 |
| 138 | Ralstonia phage RPZH6 | NA | Ralstonia pseudosolanacearum | Ralstonia phage RPZH6 | Gervaisevirus | NA | GCA_013375105.1 | ASM1337510v1 |
| 139 | Ralstonia phage RS138 | NA | Ralstonia solanacearum | Ralstonia phage RS138 | NA | NA | NC_029107 | GCA_001551265.1 |
| 140 | Ralstonia phage RSA1 | NA | Ralstonia solanacearum | Aresaunavirus RSA1 | Aresaunavirus | Peduoviridae | NC_009382 | GCA_000873125.1 |
| 141 | Ralstonia phage RSB1 | NA | Ralstonia solanacearum | Higashivirus RSB1 | Higashivirus | Autographiviridae | NC_011201 | GCA_000883015.1 |
| 142 | Ralstonia phage RSB2 | NA | Ralstonia solanacearum | Kelmasvirus RSB2 | Kelmasvirus | Autographiviridae | GCA_000918515.1 | ViralProj240595 |
| 143 | Ralstonia phage RSB3 | NA | Ralstonia solanacearum | Jiaoyazivirus RSB3 | Jiaoyazivirus | Autographiviridae | GCA_000911455.1 | ViralProj229898 |
| 144 | Ralstonia phage RSF1 | NA | Ralstonia solanacearum | Chiangmaivirus RSF1 | Chiangmaivirus | NA | GCA_001500875.1 | ViralProj307806 |
| 145 | Ralstonia phage RSJ2 | Thailand | Ralstonia solanacearum | Risjevirus RSJ2 | Risjevirus | Autographiviridae | GCA_001503995.1 | ViralProj307866 |
| 146 | Ralstonia phage RSJ5 | Thailand | Ralstonia solanacearum | Risjevirus RSJ5 | Risjevirus | Autographiviridae | GCA_001503875.1 | ViralProj307842 |
| 147 | Ralstonia phage RSK1 | NA | Ralstonia solanacearum | Firingavirus RSK1 | Firingavirus | NA | NC_022915 | GCA_000913935.1 |
| 148 | Ralstonia phage RSL2 | NA | Ralstonia | Chiangmaivirus RSL2 | Chiangmaivirus | NA | GCA_001503055.1 | ViralProj307833 |
| 149 | Ralstonia phage RsoM1USA | USA | Ralstonia solanacearum | Aresaunavirus RsoM1USA | Aresaunavirus | Peduoviridae | GCA_006083715.1 | ASM608371v1 |
| 150 | Ralstonia phage RsoM2USA | USA | Ralstonia solanacearum | Ralstonia phage RsoM2USA | NA | NA | MG752970 | GCA_018603535.1 |
| 151 | Ralstonia phage RsoP1EGY | Egypt | Ralstonia solanacearum | Gyeongsanvirus RsoP1EGY | Gyeongsanvirus | Autographiviridae | GCA_003031025.1 | ASM303102v1 |
| 152 | Ralstonia phage RsoP1IDN | Indonesia | Ralstonia solanacearum | Higashivirus RsoP1IDN | Higashivirus | Autographiviridae | GCA_002990175.1 | ASM299017v1 |
| 153 | Ralstonia phage RSP15 | NA | Ralstonia | Ralstonia phage RSP15 | NA | Ackermannviridae | GCA_001736715.1 | ViralProj343489 |
| 154 | Ralstonia phage RS-PI-1 | NA | Ralstonia solanacearum | Ampunavirus RSPI1 | Ampunavirus | Autographiviridae | GCA_002619365.1 | ASM261936v1 |
| 155 | Ralstonia phage RS-PII-1 | NA | Ralstonia solanacearum | Sukuvirus RSPII1 | Sukuvirus | Autographiviridae | GCA_002618065.1 | ASM261806v1 |
| 156 | Ralstonia phage RSY1 | NA | Ralstonia solanacearum | Arsyunavirus RSY1 | Arsyunavirus | Peduoviridae | GCA_000924355.1 | ViralProj262488 |
| 157 | Ralstonia phage Sarlave | Reunion | Ralstonia pseudosolanacearum | Ralstonia phage Sarlave | Bakolyvirus | NA | MT740746 | GCA_014262835.1 |
| 158 | Ralstonia phage Simangalove | Reunion | Ralstonia pseudosolanacearum | Bakolyvirus simangalove | Bakolyvirus | NA | NC_054946 | GCA_014262615.1 |
| 159 | Ralstonia phage UAM5 | Colombia | Ralstonia | Ralstonia phage UAM5 | NA | NA | OV121131 | GCA_921293885.1 |
| 160 | Ralstonia phage vB_RsoP_BMB50 | China | Ralstonia solanacearum | Ralstonia phage vB_RsoP_BMB50 | Higashivirus | Autographiviridae | GCA_018403885.2 | ASM1840388v1 |
| 161 | Ralstonia phage vRsoP-WF2 | Spain | Ralstonia | Gyeongsanvirus RsoP1EGY | Gyeongsanvirus | Autographiviridae | ASM979758v1 | NA |
| 162 | Ralstonia phage vRsoP-WM2 | Spain | Ralstonia | Gyeongsanvirus RsoP1EGY | Gyeongsanvirus | Autographiviridae | ASM979761v1 | NA |
| 163 | Ralstonia phage vRsoP-WR2 | Spain | Ralstonia | Gyeongsanvirus RsoP1EGY | Gyeongsanvirus | Autographiviridae | ASM979763v1 | NA |

**Table S4.** Annotation of the 45 Ralstonia solanacearum–infecting phage genomes. Values indicate the number of CDS per genome assigned to each functional category, while colors represent the corresponding percentage. Core phage functions, adaptive functions and regulatory or hypothetical genes are distinguished.

| | | | | Core phage functions | | | | Adaptative functions | | | | | | |
|---|---|---|---|---|---|---|---|---|---|---|---|---|---|---|
| Phage | Lifestyle | Total number of CDS | Unknown functions CDS | Structural proteins CDS | Assembling & Packaging CDS | DNA synthesis / Replication CDS | Lysis cassette CDS | Regulation / Gene Expression CDS | Auxiliary Metabolic Genes (AMGs) CDS | Host Interaction CDS | Integration / recombination CDS | CDS AMGs-associated | CDS Host Interaction-associated | CDS Integration-/ recombination- associated |
| P1 | Temperate | 52 | 27 | 7 | 2 | 9 | 4 | 0 | 2 | 0 | 1 | - lipid A acylase PagP<br>- deoxynucleoside monophosphate kinase | X | - integrase |
| P2 | Virulent | 64 | 26 | 9 | 2 | 12 | 4 | 5 | 3 | 2 | 1 | - acyl-CoA N-acyltransferase<br>- CysH-PAPS reductase<br>- pectin lyase-like protein | - DarB-like antirestriction<br>- DNA methyltransferase | - transposase |
| P3 | Virulent | 64 | 26 | 9 | 2 | 12 | 4 | 5 | 3 | 2 | 1 | - acyl-CoA N-acyltransferase<br>- CysH-PAPS reductase<br>- pectin lyase-like protein | - DarB-like antirestriction<br>- DNA methyltransferase | - transposase |
| P4 | Virulent | 64 | 26 | 9 | 2 | 12 | 4 | 5 | 3 | 2 | 1 | - acyl-CoA N-acyltransferase<br>- CysH-PAPS reductase<br>- pectin lyase-like protein | - DarB-like antirestriction<br>- DNA methyltransferase | - transposase |
| P5 | Virulent | 64 | 26 | 9 | 2 | 12 | 4 | 5 | 3 | 2 | 1 | - acyl-CoA N-acyltransferase<br>- CysH-PAPS reductase<br>- pectin lyase-like protein | - DarB-like antirestriction<br>- DNA methyltransferase | - transposase |
| P6 | Virulent | 64 | 26 | 9 | 2 | 11 | 4 | 4 | 3 | 2 | 3 | - acyl-CoA N-acyltransferase<br>- CysH-PAPS reductase<br>- pectin lyase-like protein | - DarB-like antirestriction<br>- DNA methyltransferase | - transposase (X3) |
| P7 | Virulent | 68 | 25 | 9 | 2 | 16 | 4 | 4 | 3 | 2 | 3 | - acyl-CoA N-acyltransferase<br>- CysH-PAPS reductase<br>- pectin lyase-like protein | - DarB-like antirestriction<br>- DNA methyltransferase | - transposase (X3) |
| P8 | Virulent | 67 | 26 | 8 | 2 | 16 | 4 | 4 | 2 | 2 | 3 | - acyl-CoA N-acyltransferase<br>- CysH-PAPS reductase | - DarB-like antirestriction<br>- DNA methyltransferase | - transposase (X3) |
| P9 | Virulent | 64 | 26 | 9 | 2 | 12 | 4 | 5 | 3 | 2 | 1 | - acyl-CoA N-acyltransferase<br>- CysH-PAPS reductase<br>- pectin lyase-like protein | - DarB-like antirestriction<br>- DNA methyltransferase | - transposase |
| P10 | Virulent | 64 | 26 | 9 | 2 | 12 | 4 | 5 | 3 | 2 | 1 | - acyl-CoA N-acyltransferase<br>- CysH-PAPS reductase<br>- pectin lyase-like protein | - DarB-like antirestriction<br>- DNA methyltransferase | - transposase |
| P11 | Virulent | 64 | 26 | 9 | 2 | 12 | 4 | 5 | 3 | 2 | 1 | - acyl-CoA N-acyltransferase<br>- CysH-PAPS reductase<br>- pectin lyase-like protein | - DarB-like antirestriction<br>- DNA methyltransferase | - transposase |
| P12 | Virulent | 67 | 24 | 11 | 2 | 14 | 4 | 6 | 3 | 2 | 1 | - acyl-CoA N-acyltransferase<br>- CysH-PAPS reductase<br>- pectin lyase-like protein | - DarB-like antirestriction<br>- DNA methyltransferase | - transposase |
| P13 | Virulent | 73 | 44 | 9 | 5 | 9 | 1 | 2 | 2 | 0 | 1 | - esterase/lipase<br>- lysine--tRNA ligase | X | - transposase |
| P14 | Virulent | 59 | 23 | 17 | 4 | 7 | 5 | 0 | 0 | 3 | 0 | X | - DNA methyltransferase<br>- pentapeptide repeat protein<br>- immunity to superinfection membrane protein | X |
| P15 | Virulent | 59 | 23 | 17 | 4 | 7 | 5 | 0 | 0 | 3 | 0 | X | - DNA methyltransferase<br>- pentapeptide repeat protein<br>- immunity to superinfection membrane protein | X |
| P16 | Virulent | 60 | 23 | 18 | 4 | 7 | 5 | 0 | 0 | 3 | 0 | X | - DNA methyltransferase<br>- pentapeptide repeat protein<br>- immunity to superinfection membrane protein | X |
| P17 | Virulent | 59 | 23 | 17 | 4 | 7 | 5 | 0 | 0 | 3 | 0 | X | - DNA methyltransferase<br>- pentapeptide repeat protein<br>- immunity to superinfection membrane protein | X |
| P18 | Virulent | 75 | 45 | 10 | 5 | 9 | 1 | 2 | 2 | 0 | 1 | - esterase/lipase<br>- lysine--tRNA ligase | X | - transposase |
| P19 | Virulent | 74 | 45 | 9 | 5 | 9 | 1 | 2 | 2 | 0 | 1 | - esterase/lipase<br>- lysine--tRNA ligase | X | - transposase |
| P20 | Virulent | 60 | 26 | 17 | 4 | 6 | 5 | 0 | 0 | 2 | 0 | X | - pentapeptide repeat protein<br>- immunity to superinfection membrane protein | X |
| P21 | Virulent | 73 | 45 | 8 | 5 | 9 | 1 | 2 | 2 | 0 | 1 | - esterase/lipase<br>- lysine--tRNA ligase | X | - transposase |
| P22 | Virulent | 73 | 43 | 9 | 5 | 10 | 1 | 2 | 2 | 0 | 1 | - esterase/lipase<br>- lysine--tRNA ligase | X | - transposase |
| P23 | Virulent | 74 | 46 | 9 | 5 | 9 | 1 | 1 | 2 | 0 | 1 | - esterase/lipase<br>- lysine--tRNA ligase | X | - transposase |
| P24 | Virulent | 73 | 44 | 9 | 5 | 9 | 1 | 2 | 2 | 0 | 1 | - esterase/lipase<br>- lysine--tRNA ligase | X | - transposase |
| P67 | Virulent | 54 | 28 | 10 | 2 | 8 | 6 | 0 | 0 | 0 | 0 | X | X | X |
| P68 | Temperate | 72 | 35 | 11 | 2 | 8 | 2 | 5 | 2 | 5 | 2 | - acyl-CoA N-acyltransferase<br>- CysH-PAPS reductase | - DarB-like antirestriction (X2)<br>- DNA methyltransferase (X2)<br>- SAM-dependent methyltransferase | - integrase, site-specific tyrosine recombinase<br>- excisionase |
| P69 | Temperate | 71 | 35 | 11 | 2 | 8 | 2 | 5 | 2 | 4 | 2 | - acyl-CoA N-acyltransferase<br>- CysH-PAPS reductase | - DarB-like antirestriction<br>- DNA methyltransferase (X2)<br>- SAM-dependent methyltransferase | - integrase, site-specific tyrosine recombinase<br>- excisionase |
| P70 | Temperate | 71 | 31 | 12 | 2 | 11 | 3 | 4 | 2 | 4 | 2 | - acyl-CoA N-acyltransferase<br>- CysH-PAPS reductase | - DarB-like antirestriction<br>- DNA methyltransferase (X2)<br>- SAM-dependent methyltransferase | - integrase, site-specific tyrosine recombinase<br>- excisionase |
| P71 | Virulent | 66 | 23 | 11 | 2 | 14 | 4 | 6 | 3 | 2 | 1 | - acyl-CoA N-acyltransferase<br>- CysH-PAPS reductase<br>- pectin lyase-like protein | - DarB-like antirestriction<br>- DNA methyltransferase | - transposase |
| P72 | Virulent | 66 | 23 | 11 | 2 | 14 | 4 | 6 | 3 | 2 | 1 | - acyl-CoA N-acyltransferase<br>- CysH-PAPS reductase<br>- pectin lyase-like protein | - DarB-like antirestriction<br>- DNA methyltransferase | - transposase |
| P73 | Virulent | 74 | 44 | 9 | 5 | 10 | 1 | 2 | 2 | 0 | 1 | - esterase/lipase<br>- lysine--tRNA ligase | X | - transposase |
| P74 | Virulent | 66 | 23 | 11 | 2 | 14 | 4 | 6 | 3 | 2 | 1 | - acyl-CoA N-acyltransferase<br>- CysH-PAPS reductase<br>- pectin lyase-like protein | - DarB-like antirestriction<br>- DNA methyltransferase | - transposase |
| P75 | Virulent | 75 | 45 | 10 | 5 | 9 | 1 | 2 | 2 | 0 | 1 | - esterase/lipase<br>- lysine--tRNA ligase | X | - transposase |
| P76 | Temperate | 48 | 20 | 6 | 3 | 8 | 5 | 1 | 4 | 0 | 1 | - deoxynucleoside monophosphate kinase<br>- Iron(III) dicitrate outer membrane transporter protein<br>- nucleoside triphosphate pyrophosphohydrolase<br>- phosphoribosyl transferase | X | - shufflon-specific DNA recombinase |
| P77 | Virulent | 67 | 28 | 9 | 2 | 12 | 4 | 5 | 3 | 2 | 2 | - acyl-CoA N-acyltransferase<br>- CysH-PAPS reductase<br>- pectin lyase-like protein | - DarB-like antirestriction<br>- DNA methyltransferase | - transposase (X2) |
| P78 | Virulent | 67 | 28 | 9 | 2 | 12 | 4 | 5 | 3 | 2 | 2 | - acyl-CoA N-acyltransferase<br>- CysH-PAPS reductase<br>- pectin lyase-like protein | - DarB-like antirestriction<br>- DNA methyltransferase | - transposase (X2) |
| P79 | Virulent | 67 | 28 | 9 | 2 | 12 | 4 | 5 | 3 | 2 | 2 | - acyl-CoA N-acyltransferase<br>- CysH-PAPS reductase<br>- pectin lyase-like protein | - DarB-like antirestriction<br>- DNA methyltransferase | - transposase (X2) |
| P80 | Virulent | 67 | 28 | 9 | 2 | 12 | 4 | 5 | 3 | 2 | 2 | - acyl-CoA N-acyltransferase<br>- CysH-PAPS reductase<br>- pectin lyase-like protein | - DarB-like antirestriction<br>- DNA methyltransferase | - transposase (X2) |
| P81 | Virulent | 73 | 44 | 9 | 5 | 9 | 1 | 2 | 2 | 0 | 1 | - esterase/lipase<br>- lysine--tRNA ligase | X | - transposase |
| P82 | Virulent | 74 | 45 | 9 | 5 | 9 | 1 | 2 | 2 | 0 | 1 | - esterase/lipase<br>- lysine--tRNA ligase | X | - transposase |
| P83 | Virulent | 74 | 45 | 9 | 5 | 9 | 1 | 2 | 2 | 0 | 1 | - esterase/lipase<br>- lysine--tRNA ligase | X | - transposase |
| P84 | Virulent | 74 | 46 | 9 | 5 | 9 | 1 | 1 | 2 | 0 | 1 | - esterase/lipase<br>- lysine--tRNA ligase | X | - transposase |
| P85 | Temperate | 50 | 22 | 8 | 3 | 8 | 3 | 1 | 4 | 0 | 1 | - deoxynucleoside monophosphate kinase<br>- Iron(III) dicitrate outer membrane transporter protein<br>- nucleoside triphosphate pyrophosphohydrolase<br>- phosphoribosyl transferase | X | - integrase |
| P86 | Temperate | 50 | 22 | 8 | 3 | 8 | 3 | 1 | 4 | 0 | 1 | - deoxynucleoside monophosphate kinase<br>- Iron(III) dicitrate outer membrane transporter protein<br>- nucleoside triphosphate pyrophosphohydrolase<br>- phosphoribosyl transferase | X | - integrase |
| P87 | Temperate | 46 | 22 | 8 | 2 | 8 | 4 | 0 | 1 | 0 | 1 | - deoxynucleoside monophosphate kinase | X | - shufflon-specific DNA recombinase |
